# The amplitude of fNIRS hemodynamic response in the visual cortex unmasks autistic traits in typically developing children

**DOI:** 10.1101/2021.07.19.452678

**Authors:** Raffaele Mazziotti, Elena Scaffei, Eugenia Conti, Viviana Marchi, Riccardo Rizzi, Giovanni Cioni, Roberta Battini, Laura Baroncelli

## Abstract

Autistic traits represent a continuum dimension across the population, with autism spectrum disorder (ASD) being the extreme end of the distribution. Accumulating evidence shows that neuroanatomical and neurofunctional profiles described in relatives of ASD individuals reflect an intermediate neurobiological pattern between the clinical population and healthy controls. This suggests that quantitative measures detecting autistic traits in the general population represent potential candidates for the development of biomarkers identifying early pathophysiological processes associated with ASD. Functional near-infrared spectroscopy (fNIRS) has been extensively employed to investigate neural development and function. In contrast, the potential of fNIRS to define reliable biomarkers of brain activity has been barely explored. Features of non-invasiveness, portability, ease of administration and low-operating costs make fNIRS a suitable instrument to assess brain function for differential diagnosis, follow-up, analysis of treatment outcomes and personalized medicine in several neurological conditions. Here, we introduce a novel standardized procedure with high entertaining value to measure hemodynamic responses (HDR) in the occipital cortex of adult subjects and children. We found that the variability of evoked HDR correlates with the autistic traits of children, assessed by the Autism-Spectrum Quotient. Interestingly, HDR amplitude was especially linked to social and communication features, representing the core symptoms of ASD. These findings establish a quick and easy strategy for measuring visually-evoked cortical activity with fNIRS that optimize the compliance of young subjects, setting the background for testing the diagnostic value of fNIRS visual measurements in the ASD clinical population.

## Introduction

Autism spectrum disorder (ASD) is a heterogeneous developmental condition that involves persistent challenges in social interactions, restricted/repetitive behaviors, and the lack of behavioral and cognitive flexibility^1^. Since the pioneering work by Lorna Wing^2^, increasing epidemiological evidence indicates that autistic traits are continuously distributed across the general population^3,4^. This is due to the complex genetic and epigenetic inheritance pattern of ASD, where multiple candidate loci contribute to the pathogenesis of the disease^5,6^. Milder autistic traits have been termed the extended or broader autism phenotype (BAP), with BAP features being particularly prevalent in first- and second-degree relatives of individuals with ASD^6–9^. Since ASD and broader autistic manifestations share common genetic variants and neurobiological susceptibility factors^10^, the general population emerges as a suitable testing bed for the development of quantitative measures detecting neurostructural and neurofunctional hallmarks of autism.

Over the last decade, the biological dimension of ASD has been largely explored, thanks to the growing availability of advanced tools to explore brain correlates of neurological disorders, including high-density EEG, magnetoencephalography, positron emission tomography, magnetic resonance imaging (MRI), and functional near-infrared spectroscopy (fNIRS)^8,11,12^. A number of studies reported defective neuroanatomical and neurofunctional features in individuals with ASD, suggesting that a dysfunction of specific brain areas might underlie the core symptoms of ASD^13–17^. Interestingly, similar neurological profiles describe the relatives of autistic probands^8^.

FNIRS is an optical imaging technique that allows quantifying oxygen consumption in different regions of the cerebral cortex, providing an indirect measure of neuronal activity^18,19^. This blood-oxygen-level-dependent (BOLD) signal is similar to that detected with functional MRI (fMRI)^20^. However, fNIRS is more tolerant to motion artifacts than fMRI, and the development of robust methods for motion detection and correction allowed to avoid sedation in children^21,22^. Furthermore, fNIRS has the advantage of being totally non-invasive, low-cost, portable, noiseless, and endowed with high experimental flexibility and no setting constraints. This methodological strength provides the fNIRS with a high ecological value for investigating neural circuit maturation either in typically developing children or clinically relevant populations^21,23^.

Although the use of fNIRS in autism research is still an emerging area, a number of studies aiming to decipher the neuronal mechanisms and circuits underlying ASD evaluated different aspects of brain function and organization, including resting-state and task-evoked responses^24,25^. Coherence analyses of resting-state hemodynamic activity showed weaker local and interhemispheric functional connectivity in different cortical regions^26–31^. Moreover, individuals on the autism spectrum present patterns of atypical activity, including reduced hemodynamic responses within specific brain regions, bilateral differences in neuronal activation and the lack of cortical specialization, in tasks ranging from sensory perception^32^ to executive functions^33^, social perception^34–37^, joint attention^38– 40^, imitation^41,42^, facial and emotional processing^43–46^, speech perception and language^47–50^. The majority of studies targeting evoked brain activity was focused on the prefrontal and the temporal cortex^25^, where symptom severity seems to be inversely correlated with the degree of cortical activation^42,43^.

Growing evidence suggests that fNIRS might be a candidate biomarker for several neuropsychiatric disorders, including ASD^29,51–55^. In particular, functional network efficiency^29^, weighted separability of NIRS signals^56^, multi-layer neural networks and sample entropy of spontaneous hemodynamic fluctuations^54,57^ have been proposed as auxiliary diagnosis indexes for ASD. However, all these approaches require complex algorithms to extract high-level features from the fNIRS raw data, while fitness, applicability and translational value of biomarkers greatly depend on their ease of use. In this framework, the analysis of visual phenotype has become an important model to evaluate cortical processing in different neurodevelopmental conditions^58–63^. Indeed, electrophysiological measurement of visually evoked responses has been introduced as a quantitative method to assess brain function in Rett syndrome^58,64^, and hemodynamic responses (HDR) emerged as a potential longitudinal biomarker for CDKL5 Deficiency Disorder and Creatine Transporter Deficiency in murine models^62,63^.

Since clinical studies suggested a dysregulation of sensory processing and functional connectivity in the visual cortex of ASD subjects^65–67^, we hypothesized that fNIRS visual measures could be related to broad autism dimensions in the general population. To address this issue, we developed a novel standardized procedure to assess HDR changes in the occipital cortex testing 40 randomly selected neuro-typical adults and 19 children, and measuring inter-individual differences in autistic traits, using the Autism-Spectrum Quotient (AQ^68,69^).

## Results

### An animated cartoon-based stimulus is able to evoke visual responses in the adult cortex

We measured the cortical HDR function^70^ elicited by a reversing checkerboard pattern in the adult population. In agreement with the previous literature^71,72^, we obtained a significant activation of the occipital cortex in response to different conditions of visual stimulation (Fig. 1 and Fig. S1). Grand averages across adult participants (see Table 1 for demographics) of Total Hb (THb), OxyHb (OHb) and DeoxyHb (DHb) concentration changes are plotted in Fig. 2. Using a classic mean-luminance grey screen as baseline, statistical analysis revealed a significant main effect of the checkerboard stimulus (S) with respect to the blank presentation for all HDR metrics (Radial Stimulus condition, RS, Fig. 2A).

**Table 1:**
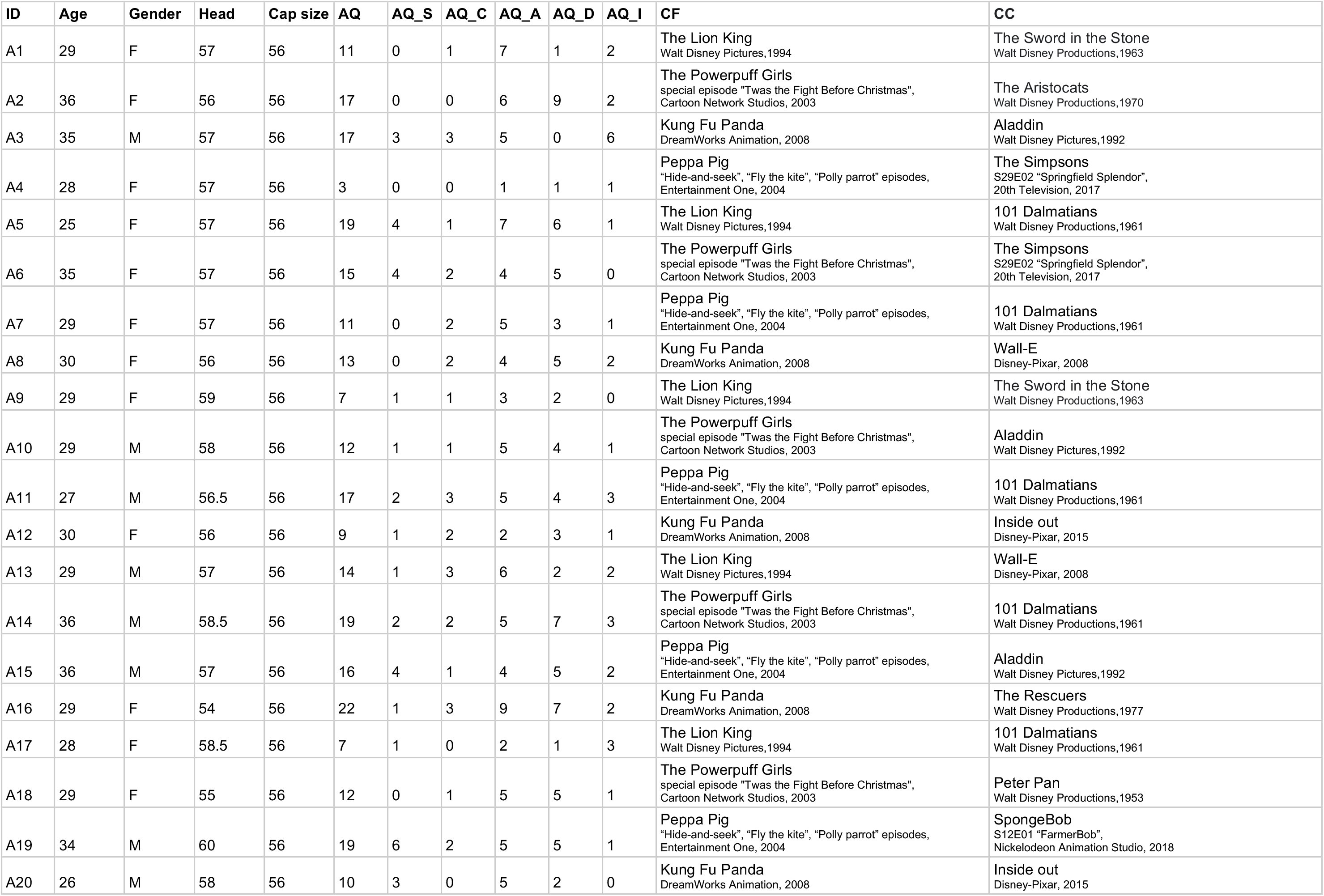
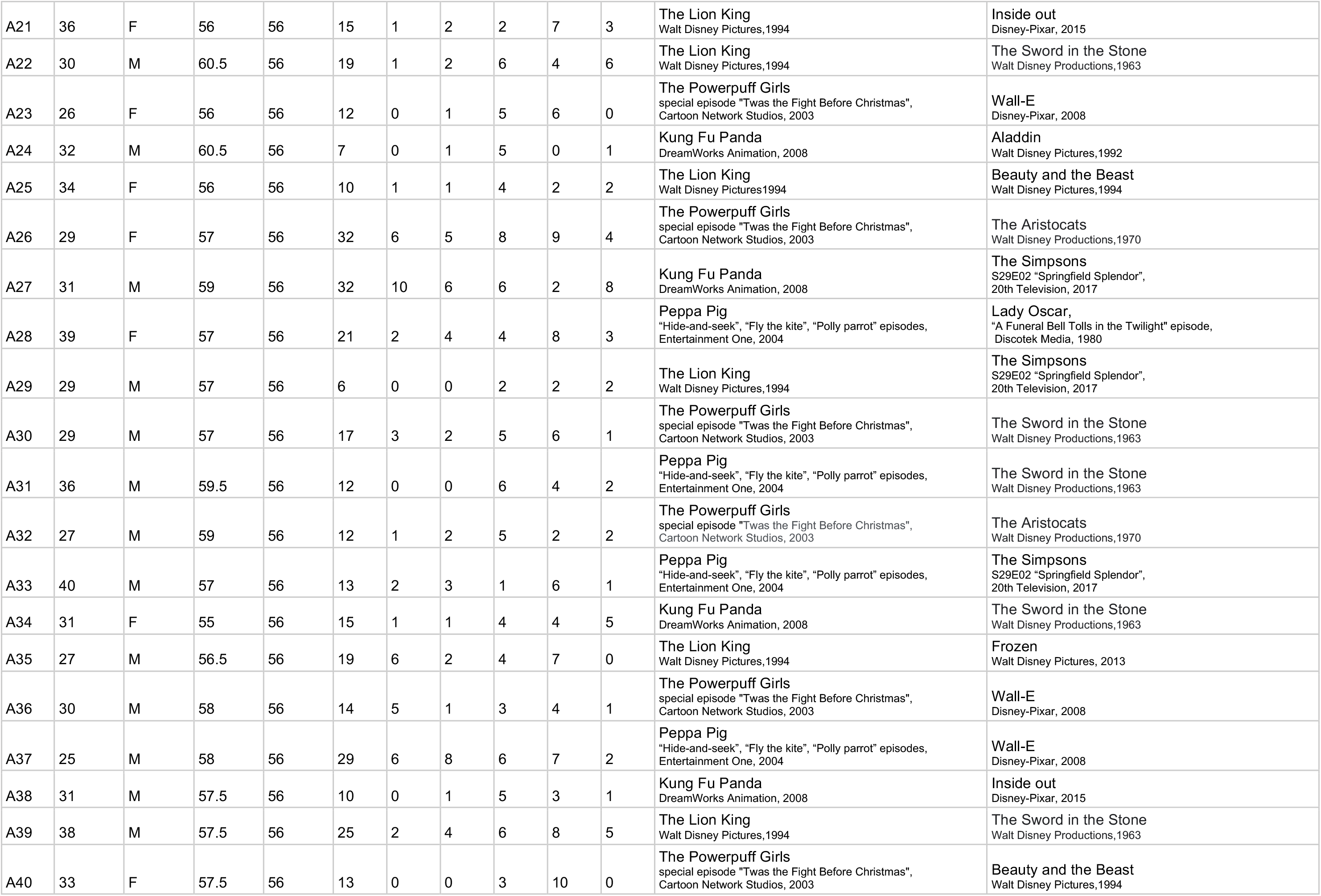
Demographic characteristics of adult subjects. Age (years), gender, head circumference (head, cm), cap size (cm), total AQ score (AQ), AQ subscale scores (AQ_S, AQ_C, AQ_A, AQ_D, AQ_I) and the movies used for visual stimulation (CF and CC according to the experimental protocol) are listed for each participant. For movies, production company, release date and episode title are indicated as well.

**Fig. 1:**
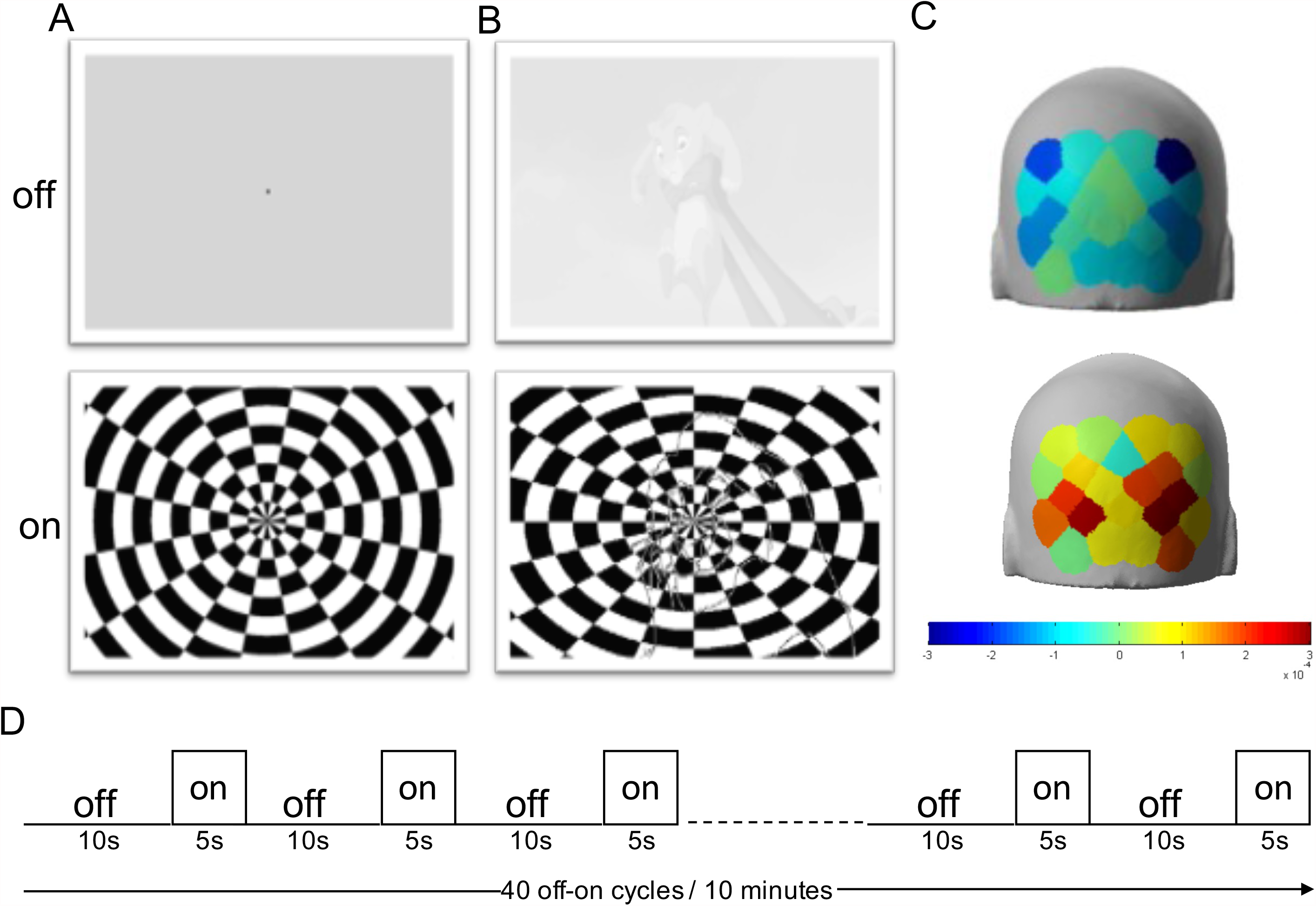
Visual stimulation and experimental paradigm. **A:** Representative frame of baseline grey screen (upper row, stimulus ‘off’) and reversing checkerboard (lower row, stimulus ‘on’) for RS condition. The small black square indicates the fixation point. **B:** Representative frame of low-contrast (20%) grey-scale baseline animated cartoon (upper row, stimulus ‘off’) and blended checkerboard-cartoon (lower row, stimulus ‘on’) for CF and CC conditions. **C:** Representative HDR in the occipital cortex during the stimulus ‘off’ (upper row) and stimulus ‘on’ activation phase (lower row) according to the output of nirsLAB software. The Look Up table is reported under the images. **D:** Experimental protocol showing that the cycles of visual stimulation were structured in blocks of 40 trials (20 trials with the reversing checkerboard and 20 trials with the ‘mock’ stimulus) for a total duration of 10 minutes.

**Fig. 2:**
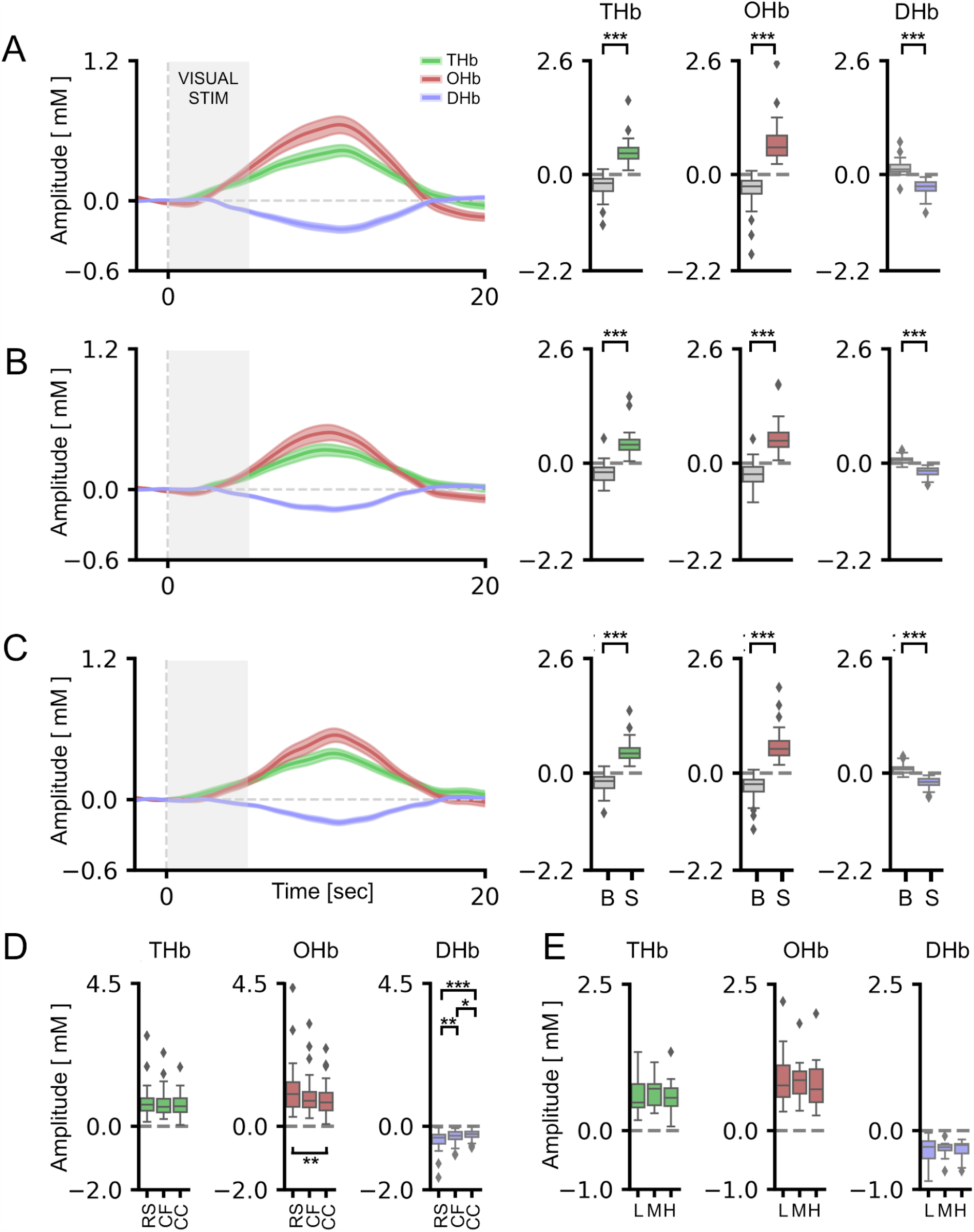
HDR was reliably detected in adults using both RS and blended RS-animated cartoons. For all panels, values in the y-axis are multiplied for 10^4. **A:** On the left, the average time course for THb (green line), OHb (red line) and DHb (blue line) in response to the RS are shown. The three plots on the right depict the average peak response to the stimulus (S) vs. the blank (B) across all the adult subjects. The stimulus-driven signal was significantly different from the blank for all the conditions (t-test, p < 0.001 for all comparisons). **B:** Same plots as above for the CF condition. On the left, the average time course of the evoked HDR is depicted. On the right, the graphs showed that the HDR amplitude was significantly higher in response to S with respect to B for THb, OHb and DHb (t-test, p < 0.001 for all comparisons). **C:** CC condition. Also in this case the S elicited significantly higher responses for THb, OHb and DHb with respect to the B (t-test, p < 0.001 for all comparisons). **D:** Comparison among different visual stimulations (RS: Radial Stimulus, CF: fixed cartoon, CC: chosen cartoon) shows no differences in evoked amplitudes for THb, whereas a significant difference was detected between RS and CC for OHb (One-way RM ANOVA, p < 0.01, post hoc BH-FDR, RS vs. CC p < 0.01) and a more complex pattern of differences emerged for DHb (One-way RM ANOVA, p < 0.001, post hoc BH-FDR, RS vs. CF p < 0.01, RS vs. CC p< 0.001, CF vs CC p < 0.05). **E:** No differences of evoked responses were detected with different contrast levels of the baseline movie (L: low, M: medium, H: high). For statistical metrics and details, refer to table S1. Data are shown as average ± s.e.m. * p < 0.05; ** p < 0.01; *** p < 0.001.

To increase the entertaining quality of our experimental paradigm, we devised an innovative visual stimulation protocol blending the checkerboard pattern with an isoluminant commercial cartoon, thus serving as a reference baseline (Fig. 1 and Fig. S1). We found a significant increase of THb and OHb, with a parallel reduction of DHb concentration, in response to S appearance, reflecting the functional activation of visual areas in this condition as well. The cortical response was independent from the cartoon employed as baseline: a comparable HDR, indeed, was clearly elicited both when the baseline movie was fixed a priori by the experimenter (Cartoon Fixed condition, CF; Fig. 2B) and when the cartoon was freely selected by the tested subject (Cartoon Chosen condition, CC; Fig. 2C). Interestingly, a significant pattern of correlations emerged among HDR metrics recorded with different stimulating conditions (Fig. S2), indicating that the quality of visual input does not quantitatively impact HDR. The amplitude of cortical activation was only slightly smaller in response to CF and CC, with the range of OHb and DHb fluctuations being significantly lower with respect to that evoked by RS (Fig. 2D).

Within the CF condition, we also established that the baseline cartoon does not affect the degree of visual activation: indeed, a comparable modification of THb, OHb and DHb concentrations was recorded using “The Lion King”, “The Powerpuff Girls”, “Peppa Pig” or “Kung Fu Panda” (Fig. S3A). Furthermore, no differences of visually evoked responses were detected modulating the contrast level of the baseline cartoon: THb, OHb and DHb fluctuations, indeed, were comparable when a fixed baseline cartoon was presented at 20%, 40%, or 80% of contrast (Fig. 2E; Fig. S3B-D). Finally, the response latency was homogenous in RS, CF, and CC conditions (Fig. S4A).

Altogether, these results demonstrate the validity of this innovative stimulation procedure to evoke a significant and reliable response in the occipital cortex preserving inter-subject variability.

### The cartoon paradigm was reliable in eliciting cortical responses in children

We measured cortical responses in typically developing children (see Table 2 for demographics) viewing the radial checkerboard blended with the animated cartoon. We compared three different conditions: each subject, indeed, was asked to select two cartoons of their preference for the baseline and the first choice was employed for the low-contrast (cartoon 1 low contrast, 20%, L1) and the high-contrast (cartoon 1 high contrast, 80%, H1) stimulation, while the second cartoon was presented only at low-contrast (cartoon 2 low contrast, L2; Fig. S1).

**Table 2:** Demographic characteristics of children. Age (years), gender, head circumference (head, cm), cap size (cm), total AQ score (AQ), AQ subscale scores (AQ_S, AQ_C, AQ_A, AQ_D, AQ_I) and the movies used for visual stimulation (C1 and C2 according to the experimental protocol) are listed for each participant. For movies, production company, release date and episode title are indicated as well.

| ID | Age | Gender | Head | Cap size | AQ | AQ-S | AQ_C | AQ_A | AQ_D | AQ_I | C1 (L1 and H1) | C2 (L2) |
| --- | --- | --- | --- | --- | --- | --- | --- | --- | --- | --- | --- | --- |
| B1 | 5 | M | 51.5 | 52 | 21 | 1 | 4 | 7 | 5 | 4 | Uncle Grandpa<br>S3E04 "Uncle Easter",<br>Cartoon Network Studios, 2016 | Teen Titans Go! To the Movies<br>Warner Bros. Animation, 2018 |
| B2 | 5 | M | 51 | 52 | 32 | 6 | 7 | 3 | 10 | 6 | Wile E. Coyote & Road Runner<br>"Coyote falls", ""Fur of flying" and "Rabid rider" episodes,<br>Acme Corporation, 2010 | ABCs song<br>Little Baby Bum - Nursery Rhymes & Kids Songs<br>youtube channel, 2014 |
| B3 | 13 | M | 57 | 56 | 26 | 6 | 3 | 4 | 7 | 6 | The Simpsons<br>S29E02 "Springfield Splendor",<br>20th Television, 2017 | Futurama,<br>S7E01 "The Bots and the Bees",<br>Comedy Central, 2012 |
| B4 | 12 | M | 56 | 56 | 44 | 8 | 8 | 11 | 13 | 4 | The Incredibles<br>Disney-Pixar, 2004 | Wile E. Coyote & Road Runner<br>"Coyote falls", ""Fur of flying" and "Rabid rider" episodes,<br>Acme Corporation, 2010 |
| B5 | 12 | M | 56 | 56 | 25 | 1 | 4 | 10 | 0 | 10 | Big Hero 6<br>Walt Disney Pictures, 2014 | The Amazing World of Gumball<br>S01E01 "The DVD",<br>Cartoon Network Development Studio Europe, 2011 |
| B6 | 6 | F | 53 | 52 | 17 | 1 | 1 | 4 | 8 | 3 | Inside out<br>Disney-Pixar, 2015 | Floopaloo<br>S1E22 "Squirrel for a Day",<br>Marc du Pontavice, 2012 |
| B7 | 7 | M | 53 | 52 | 24 | 2 | 4 | 1 | 12 | 5 | The Chipmunks<br>"Bye, George" episode,<br>DIC Entertainment, 1989 | Floopaloo<br>S1E22 "Squirrel for a Day",<br>Marc du Pontavice, 2012 |
| B8 | 4 | F | 50.5 | 52 | 38 | 6 | 11 | 7 | 11 | 3 | Frozen<br>Walt Disney Pictures, 2013 | Bolt<br>Walt Disney Pictures, 2008 |
| B9 | 4 | M | 52 | 52 | 42 | 1 | 11 | 11 | 13 | 6 | Spider-Man: Into the Spider-Verse<br>Columbia Pictures, 2018 | Bolt<br>Walt Disney Pictures, 2008 |
| B10 | 4 | M | 51 | 52 | n.a. | n.a. | n.a. | n.a. | n.a. | n.a. | Cars<br>Walt Disney Pictures, 2006 | Thomas & Friends<br>"Diesel and the duckling" episode,<br>Mattel, 2019 |
| B11 | 9 | F | 54 | 56 | 28 | 2 | 6 | 3 | 13 | 4 | The Simpsons<br>S29E02 "Springfield Splendor",<br>20th Television, 2017 | Zig & Sharko<br>S01E28 "Moby Zig",<br>Xilam Animation, 2010 |
| B12 | 8 | M | 54.5 | 56 | 26 | 4 | 5 | 5 | 4 | 8 | Beauty and the Beast<br>Walt Disney Pictures, 1994 | Ice Age<br>Blue Sky Studios, 2002 |
| B13 | 4 | M | 53 | 56 | 44 | 11 | 9 | 8 | 9 | 7 | Finding Nemo<br>Disney-Pixar 2003 | 101 Dalmatians<br>Walt Disney Productions, 1961 |
| B14 | 6 | M | 53 | 52 | 36 | 5 | 2 | 3 | 19 | 7 | Trolls<br>DreamWorks Animation, 2016 | Curious George<br>S1E03 "Zeros to Donuts",<br>Universal Animation Studios, 2006 |
| B15 | 7 | M | 51.5 | 52 | 49 | 12 | 8 | 10 | 9 | 10 | Cars<br>Walt Disney Pictures, 2006 | Pup Academy<br>S1E02 "Tell Us About Your Human Day",<br>Air Bud Entertainment, 2019 |
| B16 | 10 | F | 55 | 56 | 48 | 9 | 9 | 10 | 10 | 10 | Frozen<br>Walt Disney Pictures, 2013 | Descendants<br>Disney Channel Original Productions, 2015 |
| B17 | 10 | M | 54 | 56 | 40 | 9 | 5 | 7 | 13 | 6 | The Simpsons<br>S29E02 "Springfield Splendor",<br>20th Television, 2017 | Uncle Grandpa<br>S3E04 "Uncle Easter",<br>Cartoon Network Studios, 2016 |
| B18 | 4 | F | 49.5 | 52 | 19 | 2 | 3 | 5 | 6 | 3 | Frozen<br>Walt Disney Pictures, 2013 | Bing<br>S1E06 "Smoothie",<br>Tandem Films e Digitales Studios, 2014 |
| B19 | 7 | M | 54 | 56 | 19 | 2 | 3 | 5 | 6 | 3 | Ranger Rob<br>S1E12 "Big Stink in Big Sky Park"<br>Nelvana Enterprises Inc, 2016 | Arex&Vastatore<br>"Casket of fear" episode<br>Arex&Vastatore YouTube Channel, 2021 |

Our data showed a significant activation of the visual cortex, with a prominent change of THb, OHb and DHb concentration in response to the S with respect to the blank for all conditions tested (Fig. 3A-C). The amplitude of elicited cortical responses was comparable following L1, H1 and L2 (Fig. 3D), proving that the HDR is independent from the cartoon narrative selected for the baseline and the contrast level of baseline presentation in children as well. Small differences were observed for response latency among L1, H1 and L2 conditions (Fig. S4B). A highly significant pattern of correlations among different HDR indexes recorded in the diverse conditions was detected (Fig. S5).

**Fig. 3:**
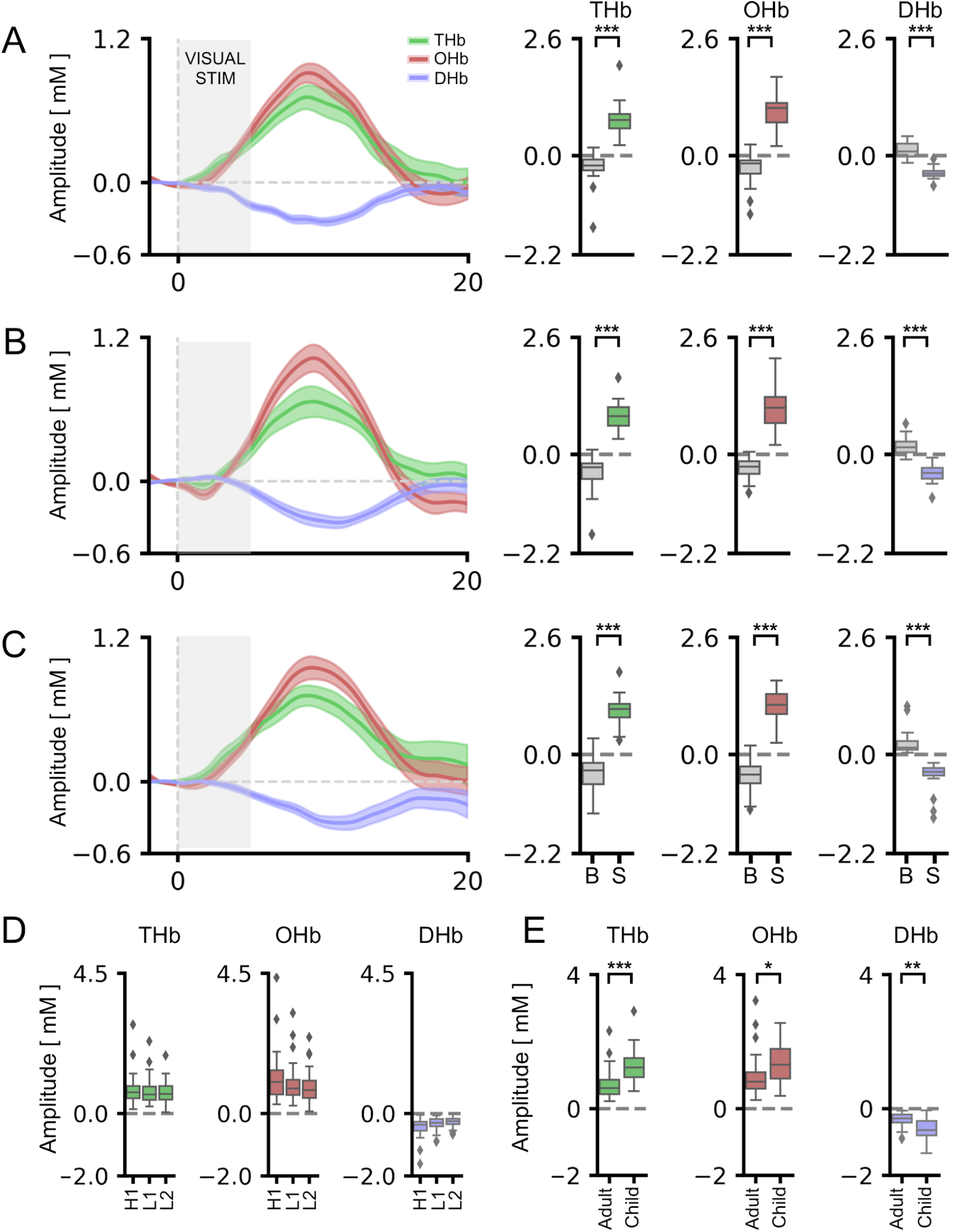
HDR signal was reliably detected in children using blended RS-animated cartoons with low and high contrast. For all panels, values in the y-axis are multiplied for 10^4. **A:** On the left, the average time course for THb (green line), OHb (red line) and DHb (blue line) in response to low-contrast (20%) blended RS-animated cartoons is shown. On the right, the graphs represent the average amplitude of the evoked HDR following the stimulus (S) and the blank (B). A significantly different response to S with respect to the B was detectable for all metrics (t-test, p < 0.001 for all comparisons). **B:** The average time course of the HDR to high-contrast (80%) blended RS-animated cartoons is shown. Here, the baseline cartoon is the same as the experiment described in panel A, but a different part of the movie was used. THb, OHb and DHb showed a significantly higher deflection to the S with respect to the B in this condition as well (t-test, p < 0.001 for all comparisons). **C:** On the left, the average time course of HDR following the second low-contrast blended RS-animated cartoon selected by the subject. On the right, the analysis of peak amplitudes revealed significantly higher responses during S compared to B for THb, OHb and DHb (t-test, p < 0.001 for all comparisons). **D:** Comparison among different contrast levels of the baseline cartoon revealed no differences in the amplitude of HDR. **E:** Response amplitudes for low-contrast blended RS-animated cartoons in adults and children. More specifically, we compared the response to CF condition of adults with L1 condition for children. The average amplitude of HDR was significantly higher in children (t-test, p < 0.001 for THb, p < 0.05 for OHb, p < 0.01 for DHb). For statistical metrics and details, refer to table S1. Data are shown as average ± s.e.m. * p < 0.05; ** p < 0.01; *** p < 0.001.

In agreement with previous literature^72^, a maturational trend of cortical responsivity was recognized, with children showing significantly higher HDR amplitude with respect to adult subjects (Fig. 3E). On the contrary, age-dependent effects were not identified for response latency (Fig. S4C).

These findings establish a novel method for measuring visually-evoked cortical activity with fNIRS that ensures an elevated compliance of young subjects and high-quality reliability of measurements, suggesting a valuable tool for studying visual cortical processing in typically developing children, but also in clinically relevant populations.

### Negative correlation of HDR amplitude with AQ score

Despite no effects detectable in adults (Fig. 4A-C), the amplitude of visual responses was highly correlated to AQ scores in children (Fig. 4D-E). Consistent with a recent work^54^, the correlation was specific for THb, with higher AQ score being associated with a lower amplitude of THb visually-evoked signals (Fig. 4D-E).

**Fig. 4:**
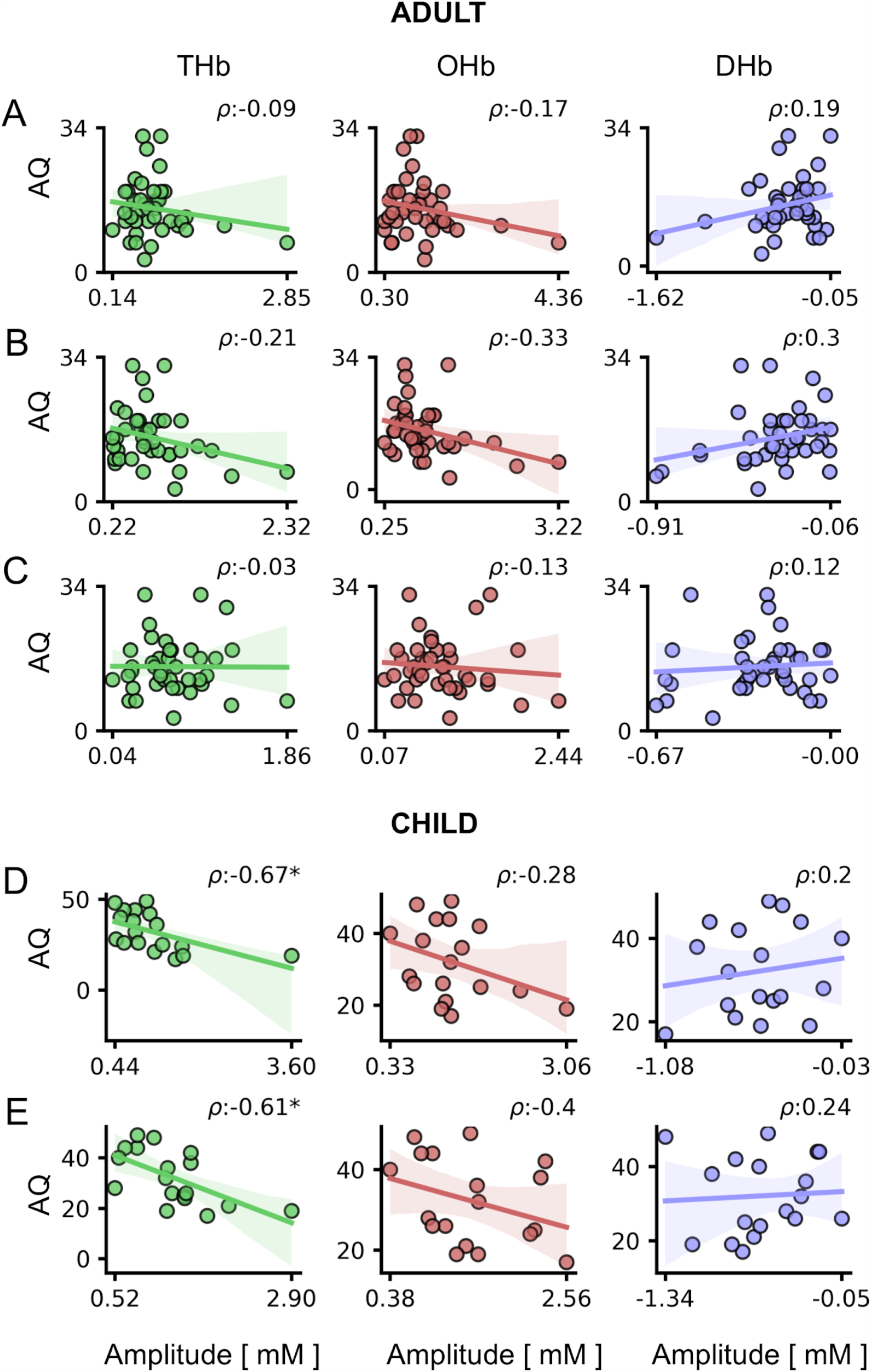
Correlation between HDR and AQ scores. For all panels, values in the x-axis are multiplied for 10^4. The ρ (rho) index in each plot indicates the Spearman correlation value. **A-C:** Correlation between HDR and AQ scores in adults, for amplitudes obtained using RS (**A**), CF (**B**), and CC (**C**). No significant correlations were detected for adult participants. **D-E:** Correlation between HDR and AQ scores in children, for amplitudes obtained using high (**D**), and low (**E**) contrast baseline cartoons. A significant correlation was found between THb and AQ scores for both high- and low-contrast blended stimuli (p < 0.05 for both cases). Circles are individual values, lines represent the linear regression model fit and shaded regions are the 95% CI.

Interestingly, HDR amplitude was especially linked to social and communication autistic traits (Fig. 5, S6): indeed, assessing separately the five AQ subscales^69^ we found a significant correlation of THb and OHb with the Social Skills subscale (AQ_S, Fig. 5A-B), while only THb modulation was related to the Communication subscale (AQ_C, Fig. 5C-D). Given the reliability across different visual tasks (L1 and H1), the strongest interaction was between THb and AQ_S (Fig. 5A-B).

**Fig. 5:**
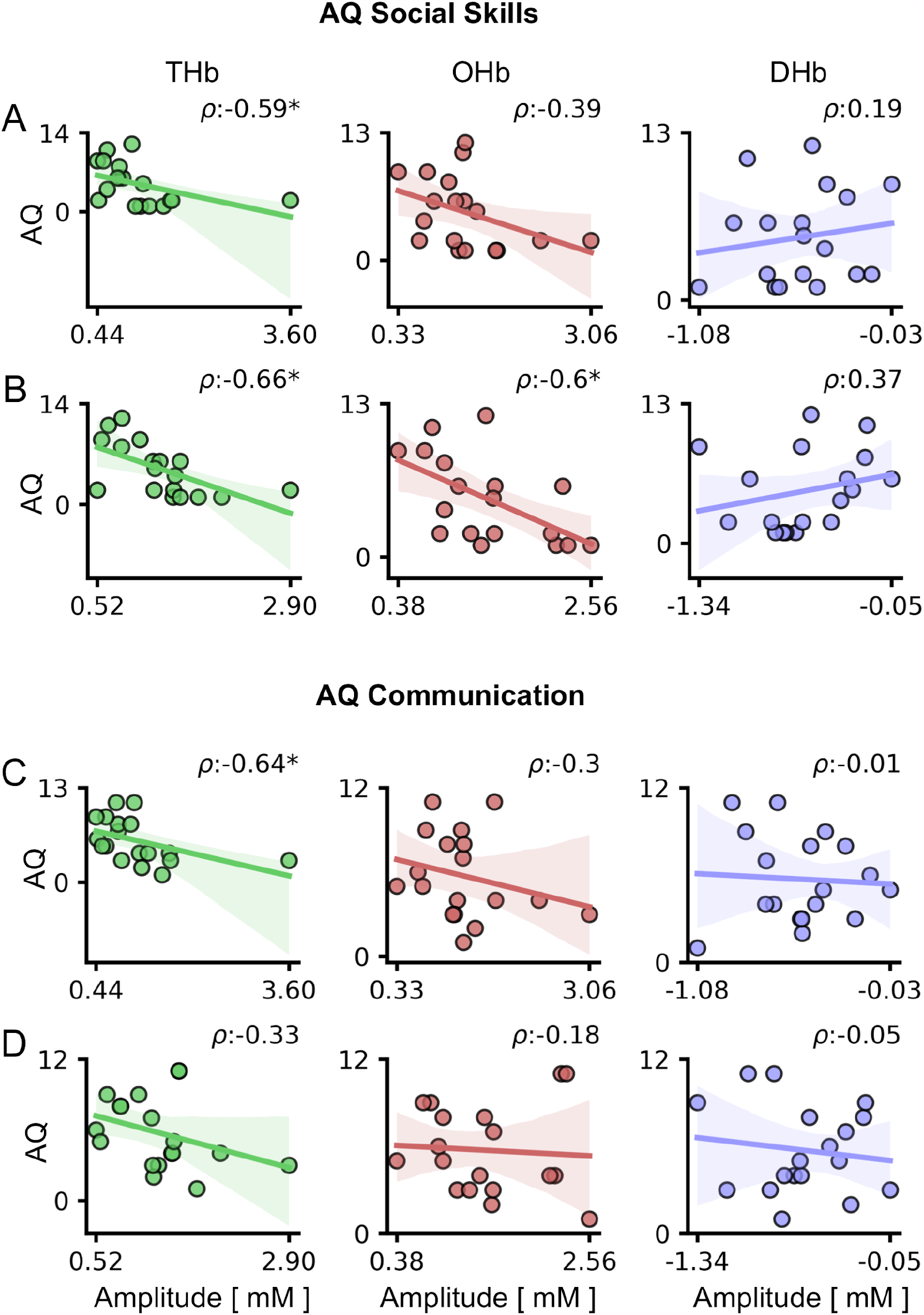
Correlation between HDR and AQ subscales in children. For all panels, values in the x-axis are multiplied for x10^4. The ρ (rho) index in each plot indicates the Spearman correlation value. **A-B:** Correlations between HDR and AQ Social Skills (AQ_S) subscale. A significant correlation between THb and AQ_S was detected using both high- (A) and low-contrast blended stimuli (B; p < 0.05 for both cases). In addition, OHb recorded in response to the low-contrast blended RS-cartoon was significantly correlated with AQ_S (B; p < 0.05). **C-D:** Correlations between HDR and AQ Communication (AQ_C) subscale. THb amplitude in response to the high-contrast blended RS-cartoon was significantly correlated with AQ_C (p < 0.05). Circles are individual values, lines represent the linear regression model fit and shaded regions are the 95% CI.

## Discussion

We measured hemodynamic responses in the occipital cortex while subjects viewed a reversing checkerboard pattern on a grey isoluminant baseline or the same stimulus blended with a commercial animated cartoon. In all participants, the patterned stimulus elicited a significant change of cortical Hb (upwards for THb and OHb, down for DHb) independently from the reference baseline, while no response was detected following blank presentation. Since HDR consists of an initial increase in oxygen rich blood followed by a smaller depletion of deoxy-haemoglobin, with a tight interrelation among total, oxygenated and reduced Hb levels, reporting all data allows for a more accurate physiological interpretation of the results^70^.

Interestingly, the level of occipital cortex activation did not depend on either the movie selected as reference baseline or the baseline contrast, with only a slight reduction of OHb and DHb change in response to the stimulus blended to animated cartoons with respect to the classic RS condition. These data demonstrate the reliability of this novel procedure with high entertaining and ecological value in eliciting cortical activity. Thus, our approach might be helpful for studying cortical function in children with an atypical trajectory of brain development, commonly showing a reduced compliance in experimental environments.

The magnitude of HDR modulation, and in particular of THb, was inversely correlated with AQ scores in children. Indeed, we found that the higher were the AQ scores of subjects the lower was the amplitude of THb response to visual stimulation, suggesting that visually-evoked fNIRS responses are able to capture the dimension of autistic traits in the general young population. Our findings are consistent with previous studies showing that cortical activation measured with fNIRS and the performance in visual psycophysics negatively reflected ASD symptom severity^42,43,73–76^. Moreover, these data reinforce the concept that THb changes could provide richer discriminative information for classifying between typically developing children and ASD subjects^54^. A tentative explanation of reduced HDR in children with stronger autistic traits might be found in the difference of perceptual styles in the general population^77,78^: the preference for focusing on local details vs. the global stimulus configuration, indeed, is a defining feature of ASD^79^ and locally centered perception could be less effective in activating neural circuits of visual cortex. Interestingly, a recent study established a vascular link to ASD, showing early dysfunction of endothelial cells and impaired endothelium-dependent vasodilation in a mouse model of 16p11.2 deletion^80^. Since the HDR measured with fNIRS strongly relies on neurovascular coupling, this suggests that lower neurophysiological activity may stem in part from endothelial-dependent vascular factors. In contrast, we failed to detect any correlation between HDR and AQ scores in adult subjects. This is likely due to the different dynamic range and amplitude of HDR responses measured in children and adult participants. Accordingly, the maturation of neural and vascular networks over the first years of life affects neurovascular coupling with a pattern of hemodynamic responses significantly different in developing and adult brain^81^.

In our experiment, the THb index describes about 45% of the variance in AQ scores. This correlation is remarkably high, considering that it is detected across two separate visual stimulating procedures, i.e., the vision of low- and the high-contrast blended RS-animated cartoons. Although the autistic questionnaire for children is well-validated, showing good test-retest reliability and high internal consistency, not all items have the same validity and factor analysis identified five subscales, named ‘Social Skills’, ‘Communication’, ‘Attention to Detail’, ‘Imagination’ and ‘Attention Switching’^69^. We observed variability in the strength of correlation between HDR and AQ subscales, with the highest correlation for the ‘Social Skills’ and ‘Communication’ subscales. Interestingly, ‘Social Skills’ and ‘Communication’ are the subscales with higher construct validity performance in differentiating individuals with or without ASD^82,83^, while ‘Attention to Detail’ was the poorest classifying domain^84^. It is also worth stressing that among the items composing the ‘Attention to Detail’ subscale, only four actually relate to sensory perception (e.g. ‘My child usually notices details that others do not’), the others being more focused on cognitive functions (e.g. ‘My child is fascinated by numbers’). Accordingly, the first pilot report of the AQ questionnaire showed lower internal consistency (measured by Cronbach’s alpha coefficient) of ‘Attention to Detail’ items compared to other subscales^68^.

Although primarily affecting social functioning, there is a growing body of evidence showing that ASD is also associated with abnormalities in multiple sensory domains, fluctuating between hyper- and hypo-sensitivity to sensory stimuli^73,85^. In addition to a higher incidence of refractive errors and strabismus^86^, anomalies in visual processing, visual attention and visual-motor integration have been described in ASD population^66,73,87^. Interestingly, sensory symptoms are correlated with the severity of the disorder, at least in children^88^. Moreover, commonly observed alterations in social skills might have a visual component^73^ and perception deficits could impact with cascading effects on the maturation of cognitive and social domains^87^.

It has been recently suggested that an early assessment of pupil size modulation and visual behavior might improve the diagnostic process of ASD^66,77,87,89^. Currently, ASD diagnosis and follow-up almost entirely rely on phenotypic information collected via clinical measures and parental input that are highly prone to subjective bias^90^. Moreover, the late appearance of some behavioral autistic traits often delays the diagnosis until mid-childhood^91,92^. Thus, the identification of solid brain biomarkers early predicting ASD pathophysiology is a critical step to anticipate tailored interventions, leading to better outcomes for patients and possibly even the prevention of certain behaviors typically associated with ASD. Objective biomarkers have also the potential to be helpful in the management of patients, allowing the classification of disease severity and monitoring response to treatments^12^. A recent systematic review highlighted that both functional and structural neuroimaging features might predict ASD diagnosis in the early pre-symptomatic period^93,94^, but further studies are needed to validate the promising performance of such biomarkers ^12^. Lately, resting-state fNIRS measurements have been suggested as candidate biomarkers for ASD^29,54,56,57^. As stressed above, fNIRS offers significant advantages with respect to other neuroimaging tools, including non-invasiveness, ease of use, no need of sedation, tolerance to movements and portability, making it a child-friendly approach. However, the extraction of metrics with diagnostic value from resting-state recordings involves complex algorithms.

In contrast, our analysis of visually evoked responses is quick, easy and requires only that children pay attention to a short movie of their choice. Since our stimulating strategy has been studied to optimize the compliance of young subjects, we believe that our results might set the background for testing the predictive value of fNIRS visual measurements to empower early detection of autistic traits. Moreover, screening for autistic traits in the general population may be helpful in epidemiological research because it may provide a large sample size to investigate the correlation between autism phenotype severity and other pathophysiological processes^83^.

## Materials and methods

### Subjects

We recruited a total of 40 adult subjects (20 women, age: 31.05 ± 3.94 (SD) years) and 19 children (5 girls, age: 7.20 ± 3.01 (SD) years). All participants reported normal or corrected-to-normal vision and had no diagnosed neuropsychiatric condition. Experimental procedures on children were authorized by the Regional Pediatrics Ethics Board (Comitato Etico Pediatrico Regionale-Azienda Ospedaliero-Universitaria Meyer-Firenze, Italy; authorization number 201/2019) and were performed according to the declaration of Helsinki. Written informed consent was obtained from all adult participants and from the parents of each child, authorizing the use of anonymized data for research purposes. Assent was also obtained from the children involved in the study before participation.

### AQ score

Adult participants filled in the Autistic-traits Quotient (AQ) questionnaire, a 50-items self-administered report validated for the Italian version^68,95^. The items consist of descriptive statements assessing personal preferences and typical behavior. For each item, participants respond on a 4-point Likert scale: “strongly agree”, “slightly agree”, “slightly disagree”, and “strongly disagree”. The items are grouped in five subscales: Social Skills, Communication, Attention to Details, Imagination and Attention Switching. All the questionnaires were scored by a neuropsychiatrist blinded to subject data: 1 point was assigned when the participant’s response was characteristic of ASD (slightly or strongly), 0 points were attributed otherwise. Total scores range between 0 and 50, with 32 being the clinical threshold for autism risk^68^. No subjects scoring above 32 points were recorded. The mean (min-max) of the scores was 15.0 (3-32) with SD of 6.5. Since child self-report might be affected by reading and comprehension difficulties, the children’s version of Autism Spectrum Quotient (Italian version of AQ-child) was completed by parents^69^. This version of the AQ questionnaire includes 50 items as well, grouped in the same subscales described above, and parents were required to report for each statement the degree of consistency with their child’s behavior. Scores range from 0 to 150, since the response scale is treated as a 4-point Likert scale with 0 representing definitely agree; 1 slightly agree; 2 slightly disagree; and 3 definitely disagree. Items were reverse scored as needed. The threshold score is 76^69^. All subjects scored below 76 points. The mean (min-max) of the scores was 32.1 (17-49) with SD of 10.7.

### Apparatus and montages

To measure changes in total Hb (THb) concentration and relative oxygenation levels (OHb and DHb) in the occipital cortex during the task, we used a continuous-wave NIRS system (NIRSport 8×8, NIRx Medical Technologies LLC, Berlin, Germany). Our NIRSport system consists of 8 red light-sources operating at 760 nm and 850 nm, and 7 detectors which can be placed into a textile EEG cap (EASYCAP, Herrsching, Germany), forming an array of 22 multi-distant channels^96^. Textile EEG caps of different sizes were used. The probe arrangement was fixed in each of the caps using grommets, optode stabilizers, colored labels and holders in order to assure comparable probe mapping over all subjects. For data recording, the Aurora Software 1.4.1.1 (NIRx Medical Technologies LLC) was employed. The sampling rate was 10.2 Hz. Visual areas were identified according to the craniocerebral topography within the international 10-20 system and the placement of the optodes was done using fOLD v2.2^97^ and NIRSite 2.0 (NIRx Medical Technologies LLC) softwares. Sources and detectors were symmetrically distributed to define 22 channels around the region of interest, each adjacent pair of sources and detectors defining one channel (min-max source-detector separation: 20-44 mm for adults, 22-30 mm for children; Fig. S2).

### Experimental Design and Visual Stimulation

Prior to the experiment, adult participants (or parents for children) filled in the AQ questionnaire. Then, subjects were asked to sit on a comfortable chair and the fNIRS cap was positioned. Optodes were placed into the cap and the calibration of light coupling between sensors and detectors was performed. All experimental sessions lasted 30 minutes. Visual stimuli were generated using Python 3 and Psychopy3 ^98^ and displayed with gamma correction on a monitor (Sharp LC-32LE352EWH, 60Hz refresh rate, 45 cd/m^2^ mean luminance, resolution of 800×600 pixels) placed 70 cm from the subject. Cortical hemodynamics in response to full-field, reversing, square wave, radial checkerboard, with abrupt phase inversion (spatial frequency: 0.33 cycles per degree, temporal frequency: 4 Hz; Fig. 1A) was evaluated in the time domain by measuring the peak-to-baseline amplitude and latency. To have an internal control with blank stimulation, we used an event-related design consisting of: i) 20 cycles of 5 seconds stimulus ‘on’ (reversing checkerboard, 90% of contrast) followed by 10 seconds stimulus ‘off’ and ii) 20 cycles of 5 seconds mock stimulus ‘on’ (reversing checkerboard, 0% of contrast) followed by 10 seconds stimulus ‘off’. The two stimulating conditions were pseudo randomly interleaved for each subject during the recording. Blocks lasted 10 minutes and participants were permitted to take rest between recordings. Fig. 1D shows a schematic representation of the experimental procedure. Visual events were synchronized with NIRSport over wireless LAN communication through the Python version of LabStreamingLayer (https://github.com/sccn/labstreaminglayer).

#### Recordings in adult participants

Experiment 1 for adults (exp1) aimed to understand whether a reliable hemodynamic signal could be recorded in response to the radial checkerboard merged with an animated cartoon. Thus, exp1 started with a 10-minutes recording using the reversing checkerboard as stimulus ‘on’ and the grey screen as stimulus ‘off’ (RS condition), and continued with the vision of two different blended animated cartoons, where the stimulus ‘on’ was a merge between the reversing checkerboard and the movie, whereas the stimulus ‘off’ was the grey-scale isoluminant cartoon (CF and CC conditions; Fig. S1). The merging procedure was achieved using Python3 OpenCV ^99^. During the appearance of the stimulus each frame was filtered using an automatic Canny edge detection algorithm (https://www.pyimagesearch.com/2015/04/06/zero-parameter-automatic-canny-edge-detection-with-python-and-opencv/), then the filtered cartoon was blended with the radial checkerboard. Each pixel of the animated cartoon with the same color of the corresponding pixel of the radial checkerboard was inverted, to obtain a fully visible image. The result was a RS with an overlayed cartoon frame (Fig 1B). The first cartoon was randomly selected by the operator within a group of 4 (“The Lion King”, “The Powerpuff Girls”, “Peppa Pig” or “Kung Fu Panda”; CF), whereas the latter was a free choice of the subject (CC). Exp2, aiming to dissect the contribution of baseline contrast to visual responses, was performed in a subset of adult participants (n = 15). Exp2 consisted of 3 consecutive recordings of a CF (“Peppa Pig”; “Hide-and-seek”, “Fly the kite”, “Polly parrot” episodes) with the modulation of the baseline contrast (20%, 40%, 80%; Fig. S1). The presentation order of different contrast levels was randomly shuffled.

#### Recordings in children

To confirm that the baseline movie and its contrast do not affect the emergence of visual responses to the radial checkerboard in children, we measured hemodynamic signals in response to 2 different blended RS-animated cartoons freely decided by the subject: cartoon 1 was presented at both low (20%, L1) and high (80%, H1) contrast, while only low contrast was recorded for cartoon 2 (L2; Fig. S1). In this case, the presentation order was decided by the child, in order to maximize subject compliance.

During the experimental sessions, data were quickly analyzed and visualized using nirsLAB software (NIRx Medical Technologies LLC, v2019.4).

### Signal Processing and Statistical analysis

Data preprocessing was completed using the Homer3 package (v1.29.8) in MATLAB (R2020a). We created a processing stream tailored on recent guidelines for analysis of fNIRS data^22^. First, the raw intensity data were converted to optical density (OD) changes (*hmR_Intensity2OD*). Then, channels showing very high or low optical intensity were excluded from further analyses using the function *hmR_PruneChannels* (dRange: 5e-04-1e+00, SNRthresh: 2; SDrange: 0.0-45.0). Motion artifacts were then removed by a multistep rejection protocol. After a step of motion artifact detection using the *hmR_MotionArtifactByChannel* function (tMotion: 1.0, tMask: 1.0; STDEVthresh 13.0; AMPthresh: 0.40), motion correction was performed with a combination of Spline interpolation (*hmR_MotionCorrectSpline*, p: 0.99) and Wavelet filtering (*hmR_MotionCorrectWavelet*, iqr: 0.80) functions^22^. The remaining uncorrected motion artifacts were identified using the *hmR_MotionArtifactByChannel*. A band-pass filter (*hmR_BandpassFilt: Bandpass_Filter_OpticalDensity*, hpf: 0.01, lpf: 0.50) was applied to decrease slow drifts and high-frequency noise, and the OD data were converted to Hb concentration changes using the modified Beer–Lambert law (*hmR_OD2Conc*, ppf: 1.0 1.0 1.0). Finally, trials of each subject were block-averaged for every stimulating condition and channel (*hmR_BlockAvg: Block_Average_on_Concentration_Data*, trange: -2.0 20.0)^22^. The resulting txt file was imported in Python as a Pandas DataFrame. For each subject, only the channel with the highest response amplitude was analyzed. The peak response was identified as the maximal value for THb and OHb and the minimum value for DHb. A grand average was taken of the 20 trials of data per stimulating condition and differences between visual stimulation ‘on’ (reversing checkerboard) and ‘off’ (blank) were compared. All data were normalised with respect to the blank-evoked response using a subtraction method. Statistical analysis was carried out using *pingouin* Python library ^100^ and the following functions: *pingouin*.*ttest* (paired and two-sided t-test), *pingouin*.*rm_anova* (one-way repeated measures ANOVA), *pingouin*.*pairwise_ttests* (post-hoc analysis), *pingouin*.*pairwise_corr* (Spearman correlation), *pingouin*.*regplot* (Linear regression). T-test, ANOVA and post-hoc analysis were used to assess differences in fNIRS peak responses following different stimulating conditions, whereas we tested the interaction between the amplitude of fNIRS measures and AQ scores with Spearman correlation and we employed the Linear regression to plot such correlation. Adjustments for multiple comparisons were performed using the Benjamini/Hochberg false discovery rate (BH-FDR) correction. The effect size calculated for the ANOVA was the generalized eta-squared. All the plots have been generated using *Matplotlib* Python library ^101^. All statistical metrics and details are reported in Table S1.

## Data availability

The datasets generated during the current study and scripts used for visual stimulation are available, respectively on Zenodo (http://doi.org/10.5281/zenodo.5101912) and GitHub website (https://github.com/raffaelemazziotti/FNIRS_code).

## Acknowledgements

This work was supported by a Telethon grant from GP19177 to LB, and a grant from University of Pisa (PRA-2020-50) to RB.

## Author contribution

RB, GC and LB conceived the study. RM, ES, EC, VM, RR and LB designed the experiments. RM, ES and LB carried out the research. RM, ES and LB analyzed the data. LB wrote the manuscript. All authors were involved in the revision of the draft manuscript and have agreed to the final content.

