## Supplementary material for "The amplitude of fNIRS hemodynamic response in the visual cortex unmasks autistic traits in typically developing children"

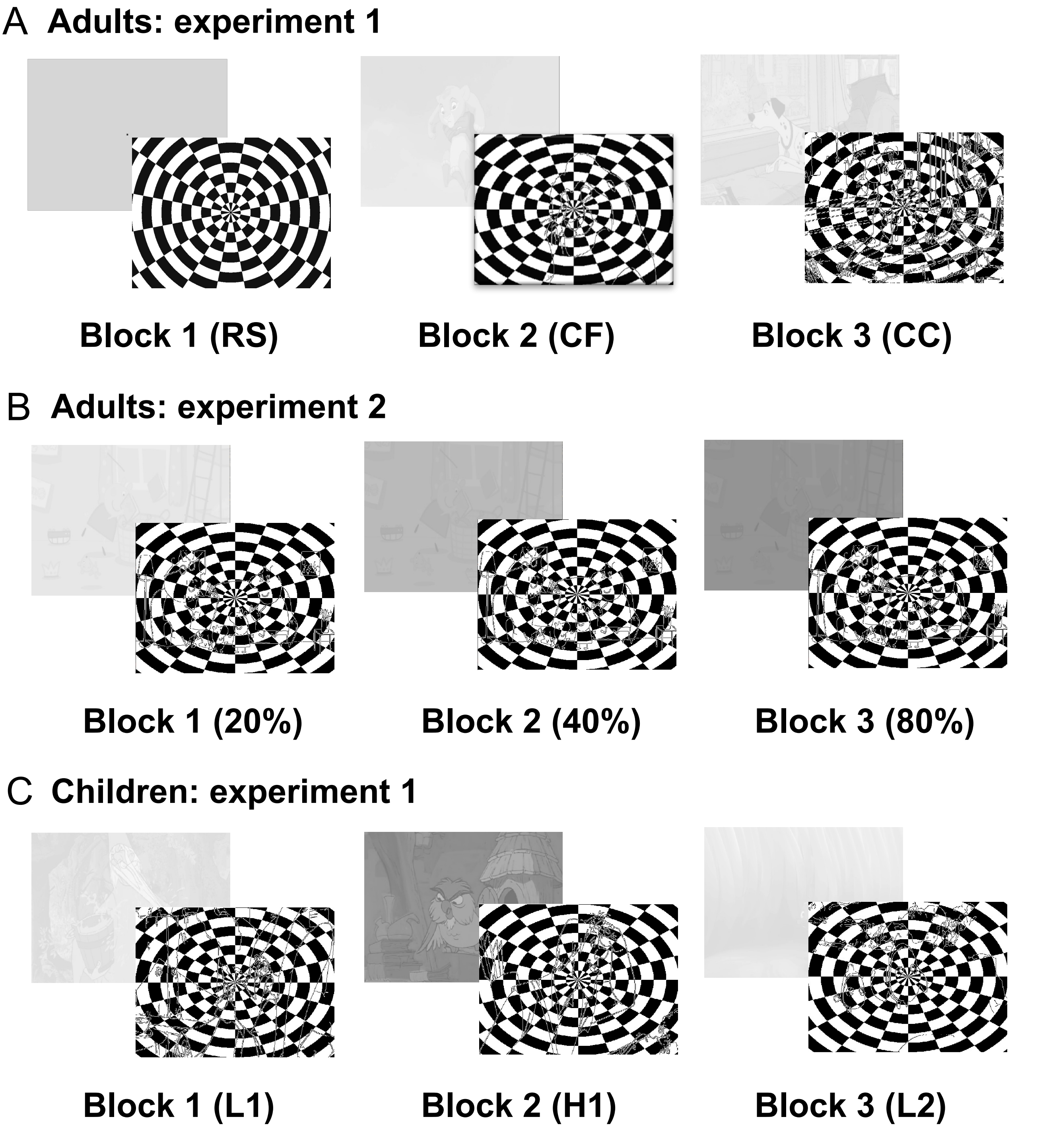

**Fig. S1:** Experimental plan. For each block (i.e., stimulating conditions), representative frames of baseline screen (stimulus 'off', up) and reversing checkerboard (stimulus 'on', low) were shown. **(A)** Experiment 1 for adult participants consisted in three 10-minute blocks, starting with the vision of the classic radial checkerboard-on-grey stimulus (RS condition) and continuing with two different RS-blended animated cartoons. The first cartoon was decided by the operator within a group of 4 (CF condition), whereas the latter was a free choice of the subject (CC condition). **(B)** Experiment 2 was carried out in a subset of adult participants (n = 15), watching the same «Peppa Pig» episodes with the modulation of the baseline contrast (20%, 40&, 80%). **(C)** Children were recorded during the vision of 2 different blended animated cartoons freely selected by the subject: cartoon 1 was shown at both low (20%, L1) and high (80%, H1) contrast, while only low contrast was used for cartoon 2 (L2).

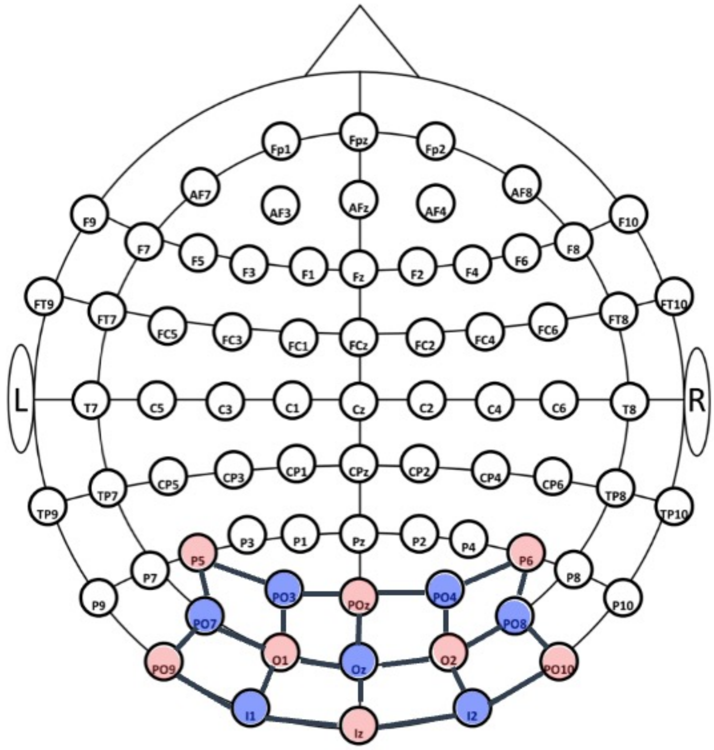

**Fig. S2:** Occipital 8x7 montage according to the international 10-20 system (the only differences in the correspondent adult occipital montage are: PPO5h for P5 and PPO6h for P6); sources (red circles) and detectors (blue circles) were symmetrically distributed to define 22 channels around the visual areas of the cortex, each adjacent pair of sources and detectors defining one channel.

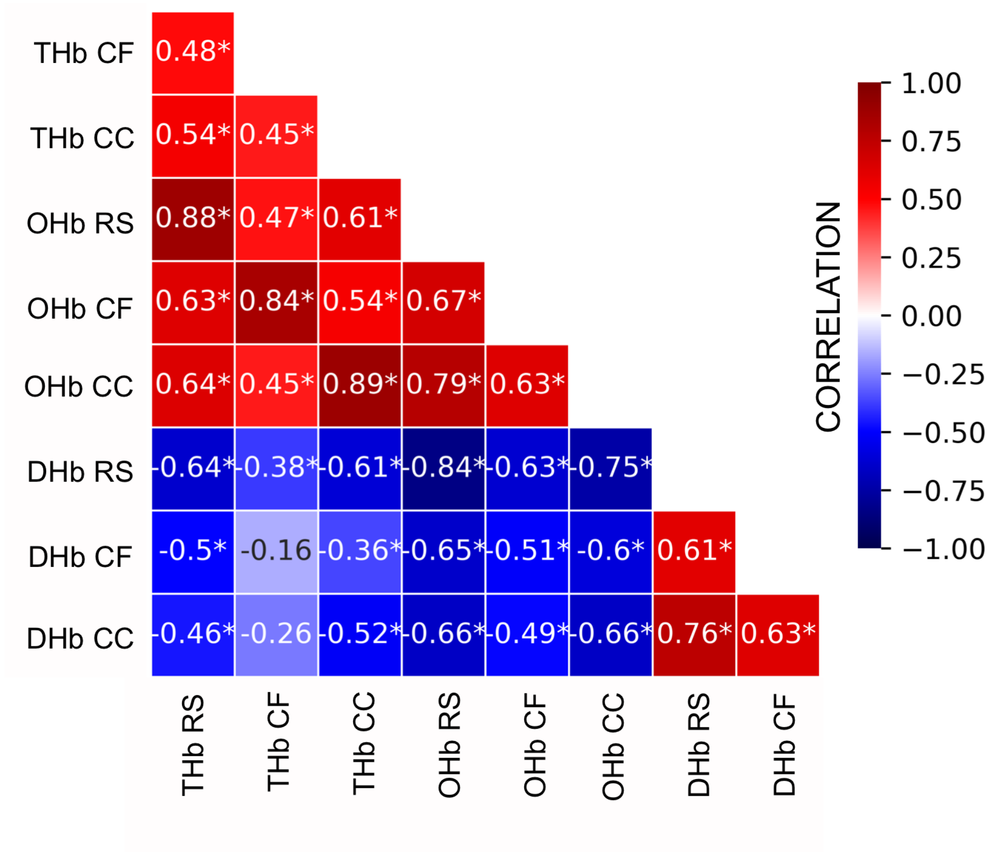

**Fig. S3: Matrix of correlations among different HDR metrics recorded in response to the diverse visual stimulations presented to adult participants.** A highly significant pattern of correlations emerged, indicating that stimulating conditions were largely interchangeable. Asterisks indicate significant correlations.

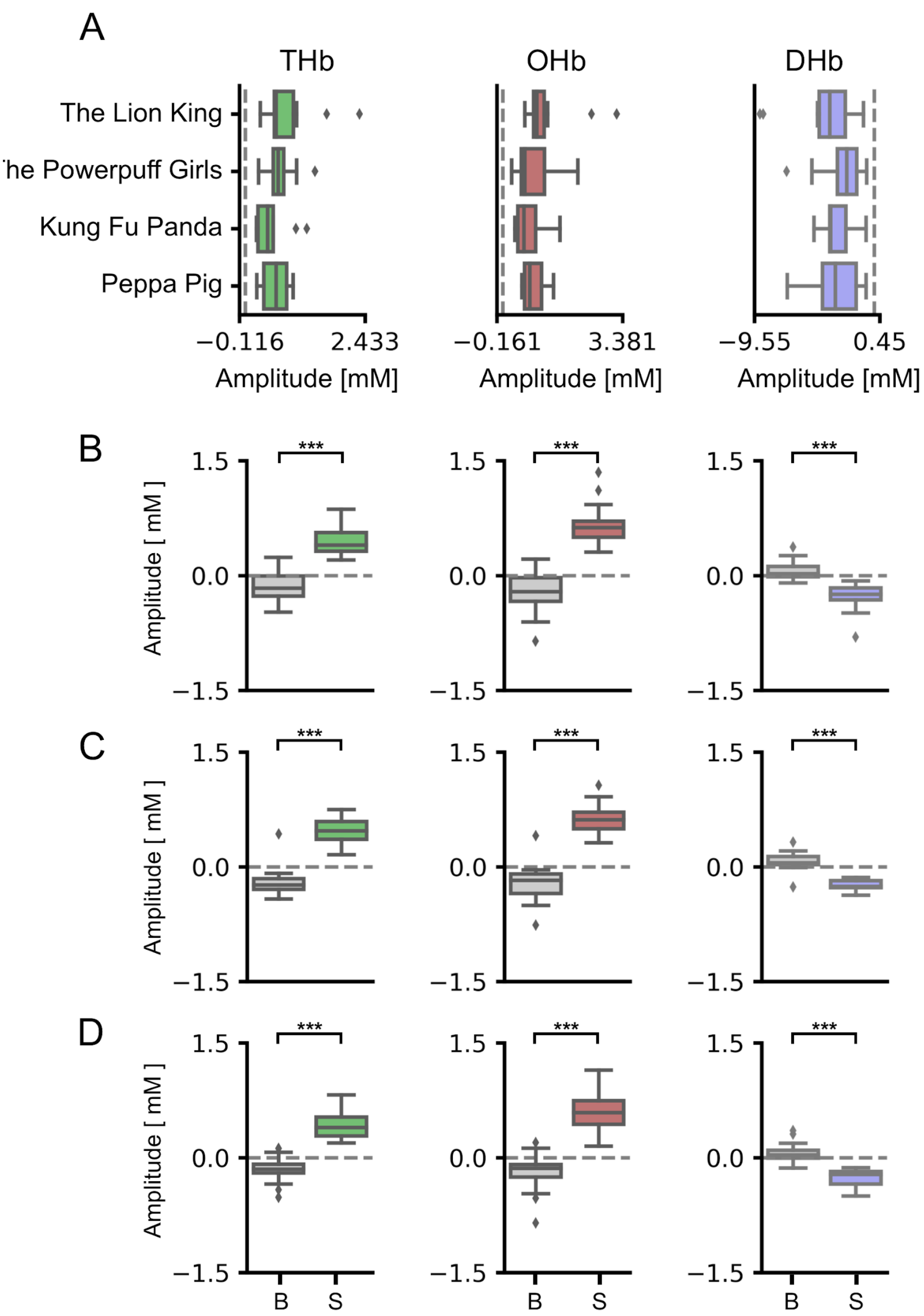

**Fig. S4: The amplitude of evoked responses did not depend on either the cartoon selected as reference baseline or the contrast level of the movie. A:** Values in the x-axis are multiplied for 10^4. No significant differences were detected in the cortical response elicited by RS blended with different baseline cartoons in adults. **B-D:** Comparison between the stimulus (S)-evoked amplitude against blank (B) using blended cartoons of low (B), medium (C) or high (D) contrast level in adults. The stimulus-evoked amplitude was significantly higher than that recorded during the B (t-test, $p < 0.001$ for all comparisons).

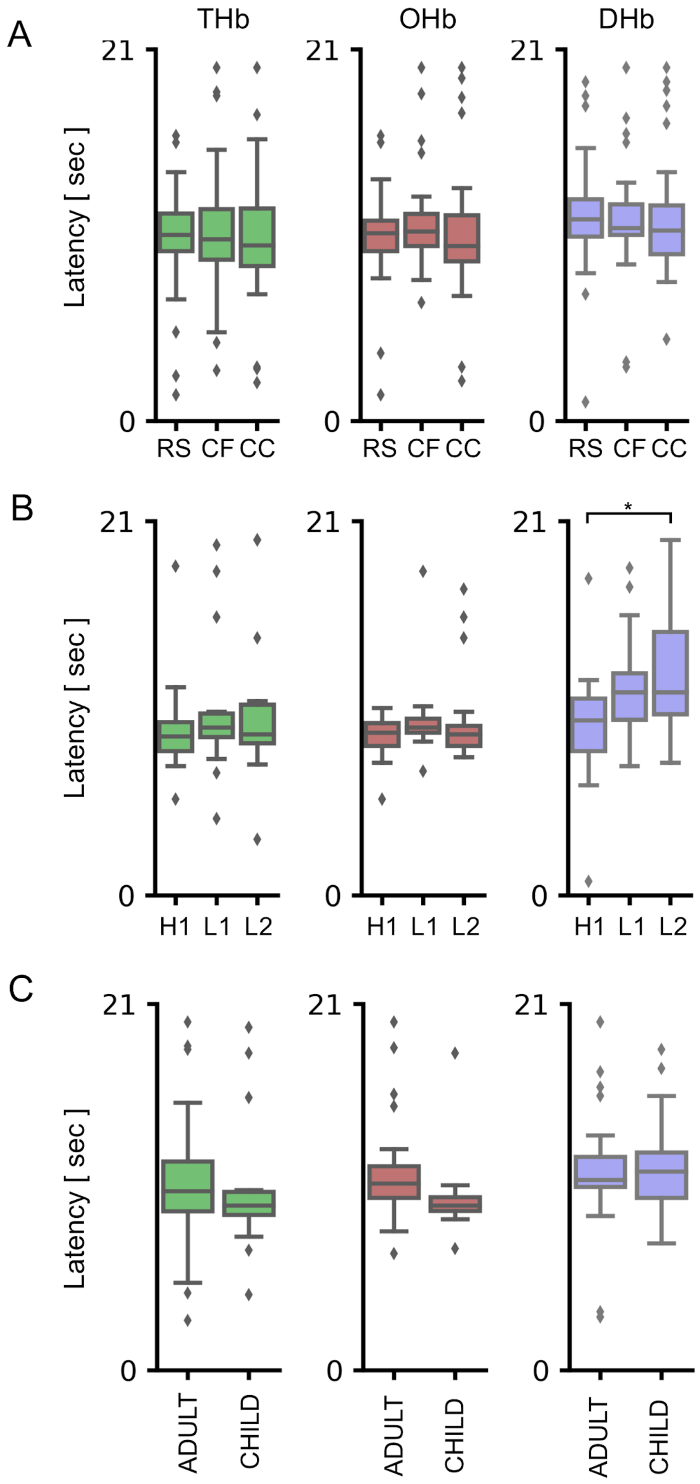

**Fig. S5: Latency of HDR in adults and children. A:** The analysis of latency to peak in adults revealed no differences among RS, CF and CC. **B:** In children, no differences of HDR latencies were found between H1 and L1, whereas a significantly higher latency of DHb was detected between H1 and L2 (One-way RM ANOVA, $p < 0.05$, post hoc BH-FDR H1 vs. L2 $p < 0.05$). **C:** No significant differences were identified for latencies between adults and children. * $p < 0.05$.

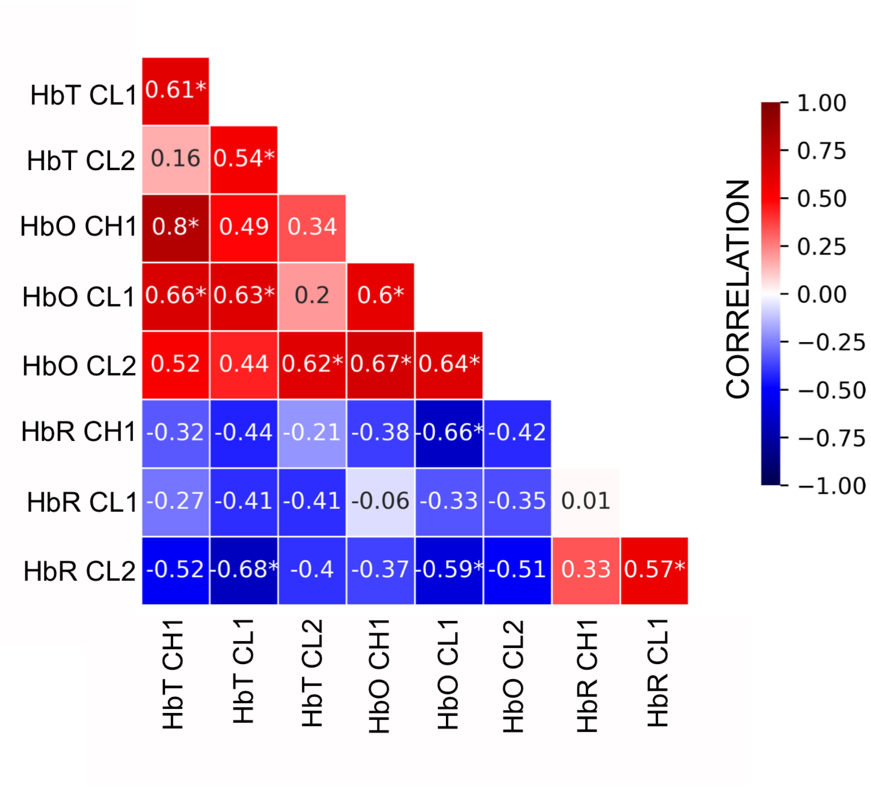

**Fig. S6: Matrix of correlations among different HDR metrics recorded in response to the diverse visual stimulations presented to children.** Similarly to the recordings in adults, a highly significant pattern of correlations was found.. Asterisks indicate significant correlations.

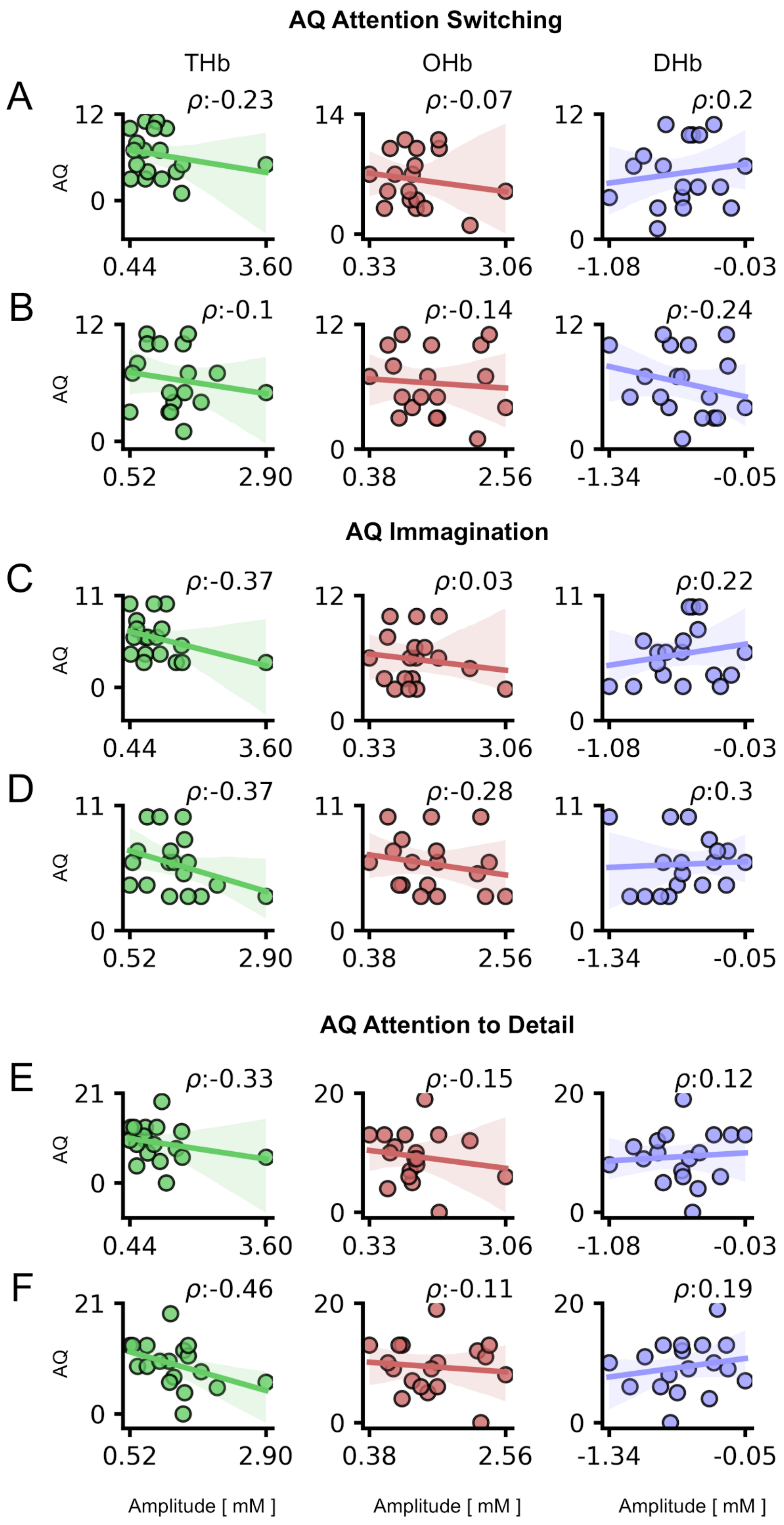

**Fig. S7: Correlation between HDR and AQ subscales in children.** For all panels, values in the x-axis are multiplied for 10^4. The ρ (rho) index in each plot indicates the Spearman correlation value. No significant correlation was present

Between THb, OHb, DHb metrics and Attention Switching (A,B), Imagination (C,D) and Attention to Detail (E,F) AQ subscales.

**Supplementary Table 1.** Statistical Table. **Hb**: Hemoglobin, **cond**: condition, **T**: t-distribution value, **power**: the achieved power of the test (1 - type II error), **BF10**: Bayes Factor of the alternative hypothesis, **p-val**: p-value (corrected for sphericity violation in anova rm and multiple comparisons in post hoc), **ng2**: generalized eta-squared, **eps**: Greenhouse-Geisser epsilon factor, **corr**: correction, i.e., multiple comparisons method, **fdr_bh**: Benjamini/Hochberg FDR correction, **r**: regression coefficient, **F**: F-statistic, **eta-square**: eta-squared effect size.

| **Figure** | **Hb** | **cond** | **test** | **T** | **tail** | **p-val** | **power** | **BF10** | | |
|---|---|---|---|---|---|---|---|---|---|---|
| 2 A | THb | rs | T-test | -10.27664192 | two-sided | 1.17E-12 | 1 | 7.09E+09 | | |
| 2 B | THb | cf | T-test | -10.53736721 | two-sided | 5.69E-13 | 1 | 1.42E+10 | | |
| 2 C | THb | cc | T-test | -11.45204453 | two-sided | 4.80E-14 | 1 | 1.52E+11 | | |
| 2 A | OHb | rs | T-test | -9.408641697 | two-sided | 1.40E-11 | 1 | 6.64E+08 | | |
| 2 B | OHb | cf | T-test | -10.10992018 | two-sided | 1.88E-12 | 1 | 4.53E+09 | | |
| 2 C | OHb | cc | T-test | -10.48477915 | two-sided | 6.58E-13 | 1 | 1.24E+10 | | |
| 2 A | DHb | rs | T-test | 9.473101868 | two-sided | 1.16E-11 | 1 | 7.94E+08 | | |
| 2 B | DHb | cf | T-test | 9.910922592 | two-sided | 3.29E-12 | 1 | 2.64E+09 | | |
| 2 C | DHb | cc | T-test | 9.799223827 | two-sided | 4.53E-12 | 1 | 1.95E+09 | | |
| **Figure** | **Hb** | **Test** | **F** | **p-val** | **ng2** | **eps** | | | | |
| 2 D | THb | anova_rm | 1.640868453 | 0.2004303395 | 0.009952442677 | 0.9901442088 | | | | |
| 2 D | OHb | anova_rm | 7.340360394 | 0.001199783266 | 0.03338550065 | 0.8892069675 | | | | |
| | | **post hoc** | **A** | **B** | **T** | **Tail** | **p-val** | **corr** | **eta-square** | **BF10** |
| | | t-test | best_cc_delta | best_cf_delta | -1.672 | two-sided | 0.102 | fdr_bh | 0.008 | 0.61 |
| | | t-test | best_cc_delta | best_rs_delta | -3.543 | two-sided | 0.003 | fdr_bh | 0.044 | 29.388 |
| | | t-test | best_cf_delta | best_rs_delta | -2.209 | two-sided | 0.05 | fdr_bh | 0.017 | 1.494 |
| **Figure** | **Hb** | **Test** | **F** | **p-val** | **ng2** | **eps** | | | | |
| 2 D | DHb | anova_rm | 16.04958967 | 1.45E-06 | 0.08357575091 | 0.8286713646 | | | | |
| | | **post hoc** | **A** | **B** | **T** | **Tail** | **p-val** | **corr** | **eta-square** | **BF10** |
| | | t-test | best_cc_delta | best_cf_delta | 2.145 | two-sided | 0.038 | fdr_bh | 0.015 | 1.329 |
| | | t-test | best_cc_delta | best_rs_delta | 5.05 | two-sided | 0 | fdr_bh | 0.102 | 1887.497 |
| | | t-test | best_cf_delta | best_rs_delta | 3.482 | two-sided | 0.002 | fdr_bh | 0.05 | 25.2 |
| **Figure** | **Hb** | **Test** | **F** | **p-val** | **ng2** | **eps** | | | | |
| 2 E | THb | anova rm | 0.690166 | 0.509822 | 0.018364 | 0.941664 | | | | |
| 2 E | OHb | anova rm | 0.672696 | 0.518386 | 0.010916 | 0.955056 | | | | |
| 2 E | DHb | anova rm | 0.14099 | 0.86911 | 0.004897 | 0.863214 | | | | |
| **Figure** | **Hb** | **cond** | **test** | **tail** | **T** | **p-val** | **power** | **BF10** | | |
| 3 A | THb | ch1 | T-test | two-sided | -6.710863 | 2.72E-06 | 1 | 7192.454 | | |
| 3 B | THb | cl1 | T-test | two-sided | -10.094777 | 7.72E-09 | 1 | 1.58E+06 | | |
| 3 C | THb | cl2 | T-test | two-sided | -8.874766 | 5.42E-08 | 1 | 2.60E+05 | | |

| | | | | | | | | |
|---|---|---|---|---|---|---|---|---|
| 3 A | OHb | ch1 | T-test | two-sided | -8.766933 | 6.50E-08 | 1 | 2.20E+05 |
| 3 B | OHb | cl1 | T-test | two-sided | -9.801022 | 1.22E-08 | 1 | 1.04E+06 |
| 3 C | OHb | cl2 | T-test | two-sided | -9.960689 | 9.49E-09 | 1 | 1.30E+06 |
| 3 A | DHb | ch1 | T-test | two-sided | 8.570421 | 9.07E-08 | 1 | 1.62E+05 |
| 3 B | DHb | cl1 | T-test | two-sided | 8.080983 | 2.12E-07 | 1 | 7.40E+04 |
| 3 C | DHb | cl2 | T-test | two-sided | 5.295974 | 4.91E-05 | 1 | 527.727 |
| **Figure** | **Hb** | **Test** | **F** | **p-val** | **ng2** | **eps** | | |
| 3 D | THb | anova_rm | 2.269052 | 0.118006 | 0.038296 | 0.6744 | | |
| 3 D | OHb | anova_rm | 2.266482 | 0.118275 | 0.034911 | 0.920718 | | |
| 3 D | DHb | anova_rm | 2.189169 | 0.126699 | 0.05409 | 0.822845 | | |
| **Figure** | **Hb** | **test** | **tail** | **T** | **p-val** | **power** | **BF10** | |
| 3 E | THb | T-test | two-sided | -4.135410352 | 0.000292627372 | 0.9944978152 | 188.71 | |
| 3 E | OHb | T-test | two-sided | -2.604585851 | 0.01352512184 | 0.7389200153 | 4.225 | |
| 3 E | DHb | T-test | two-sided | 3.536942192 | 0.001623564012 | 0.9834889618 | 36.582 | |
| **Figure** | **Y** | **X** | **method** | **tail** | **r** | **p-val** | **corr** | **power** |
| 4 A | AQ | THb_rs | spearman | two-sided | -0.08912568744 | 0.6575061582 | fdr_bh | 0.0848300996 |
| 4 B | AQ | THb_cf | spearman | two-sided | -0.2148616097 | 0.5426725338 | fdr_bh | 0.2684559699 |
| 4 C | AQ | THb_cc | spearman | two-sided | -0.028328228 | 0.8622451739 | fdr_bh | 0.0530912285 |
| 4 A | AQ | OHb_rs | spearman | two-sided | -0.1663930469 | 0.548662905 | fdr_bh | 0.1778793293 |
| 4 B | AQ | OHb_cf | spearman | two-sided | -0.3325037526 | 0.2732954373 | fdr_bh | 0.5655757921 |
| 4 C | AQ | OHb_cc | spearman | two-sided | -0.1295004709 | 0.6025285747 | fdr_bh | 0.1256529156 |
| 4 A | AQ | DHb_rs | spearman | two-sided | 0.1896391343 | 0.5426725338 | fdr_bh | 0.2182707866 |
| 4 B | AQ | DHb_cf | spearman | two-sided | 0.2991874977 | 0.2732954373 | fdr_bh | 0.4750679664 |
| 4 C | AQ | DHb_cc | spearman | two-sided | 0.117924484 | 0.6025285747 | fdr_bh | 0.1122415123 |
| **Figure** | **Y** | **X** | **method** | **tail** | **r** | **p-val** | **corr** | **power** |
| 4 E | AQ | THb_ch1 | spearman | two-sided | -0.6687346538 | 0.01446356566 | fdr_bh | 0.8925776968 |
| 4 D | AQ | THb_cl1 | spearman | two-sided | -0.6129206333 | 0.0205177254 | fdr_bh | 0.8069164533 |
| 4 E | AQ | OHb_ch1 | spearman | two-sided | -0.2801036958 | 0.3903842975 | fdr_bh | 0.2069161503 |
| 4 D | AQ | OHb_cl1 | spearman | two-sided | -0.4031012597 | 0.1943584346 | fdr_bh | 0.3952742327 |
| 4 E | AQ | DHb_ch1 | spearman | two-sided | 0.1994834439 | 0.4274269322 | fdr_bh | 0.1255757915 |
| 4 D | AQ | DHb_cl1 | spearman | two-sided | 0.2387599769 | 0.4080103079 | fdr_bh | 0.1610802525 |
| **Figure** | **Y** | **X** | **method** | **tail** | **r** | **p-val** | **corr** | **power** |
| 5 B | AQ | THb_ch1 | spearman | two-sided | -0.5917912581 | 0.01935319552 | fdr_bh | 0.7690513401 |
| 5 A | AQ | THb_cl1 | spearman | two-sided | -0.6566164489 | 0.01845705021 | fdr_bh | 0.8760130618 |
| 5 B | AQ | OHb_ch1 | spearman | two-sided | -0.3931334153 | 0.159807852 | fdr_bh | 0.3773141518 |
| 5 A | AQ | OHb_cl1 | spearman | two-sided | -0.5991102313 | 0.01935319552 | fdr_bh | 0.7824451917 |
| 5 B | AQ | DHb_ch1 | spearman | two-sided | 0.1913388697 | 0.4469126771 | fdr_bh | 0.1191550365 |

| | | | | | | | | |
|---|---|---|---|---|---|---|---|---|
| 5 A | AQ | DHb_cl1 | spearman | two-sided | 0.3669942254 | 0.1609306206 | fdr_bh | 0.3322869059 |
| **Figure** | **X** | **Y** | **method** | **tail** | **r** | **p-val** | **corr** | **power** |
| 5 D | AQ | THb_ch1 | spearman | two-sided | -0.6417570048 | 0.02454598298 | fdr_bh | 0.8540832004 |
| 5 C | AQ | THb_cl1 | spearman | two-sided | -0.3333398034 | 0.4627528962 | fdr_bh | 0.2791221623 |
| 5 D | AQ | OHb_ch1 | spearman | two-sided | -0.296994342 | 0.4627528962 | fdr_bh | 0.2282139392 |
| 5 C | AQ | OHb_cl1 | spearman | two-sided | -0.1796504237 | 0.7134812865 | fdr_bh | 0.1104870383 |
| 5 D | AQ | DHb_ch1 | spearman | two-sided | -0.008307534042 | 0.9739011846 | fdr_bh | 0.0493146419 |
| 5 C | AQ | DHb_cl1 | spearman | two-sided | -0.05192208776 | 0.9739011846 | fdr_bh | 0.0540728454 |
| **Figure** | **X** | **Y** | **method** | **tail** | **r** | **p-val** | **corr** | **power** |
| S3 | THb_rs | THb_cf | spearman | two-sided | 0.4769230769 | 0.00239365504 | fdr_bh | 0.8908005329 |
| S3 | THb_rs | THb_cc | spearman | two-sided | 0.5401500938 | 0.00052556136 | fdr_bh | 0.960373298 |
| S3 | THb_rs | OHb_rs | spearman | two-sided | 0.8771106942 | 2.07E-12 | fdr_bh | 0.9999999999 |
| S3 | THb_rs | OHb_cf | spearman | two-sided | 0.6307692308 | 2.98E-05 | fdr_bh | 0.9953861393 |
| S3 | THb_rs | OHb_cc | spearman | two-sided | 0.638836773 | 2.54E-05 | fdr_bh | 0.996387892 |
| S3 | THb_rs | DHb_rs | spearman | two-sided | -0.6399624765 | 2.54E-05 | fdr_bh | 0.996512112 |
| S3 | THb_rs | DHb_cf | spearman | two-sided | -0.4968105066 | 0.00153752136 | fdr_bh | 0.9178746454 |
| S3 | THb_rs | DHb_cc | spearman | two-sided | -0.4575984991 | 0.00358708400 | fdr_bh | 0.8596833853 |
| S3 | THb_cf | THb_cc | spearman | two-sided | 0.4508442777 | 0.00404794622 | fdr_bh | 0.8476971873 |
| S3 | THb_cf | OHb_rs | spearman | two-sided | 0.4686679174 | 0.00283870362 | fdr_bh | 0.878087982 |
| S3 | THb_cf | OHb_cf | spearman | two-sided | 0.8365853659 | 1.60E-10 | fdr_bh | 0.9999999762 |
| S3 | THb_cf | OHb_cc | spearman | two-sided | 0.4497185741 | 0.00404794622 | fdr_bh | 0.8456444659 |
| S3 | THb_cf | DHb_rs | spearman | two-sided | -0.3833020638 | 0.01596056443 | fdr_bh | 0.6997108556 |
| S3 | THb_cf | DHb_cf | spearman | two-sided | -0.1628517824 | 0.3153652463 | fdr_bh | 0.1722330029 |
| S3 | THb_cf | DHb_cc | spearman | two-sided | -0.2583489681 | 0.1105587073 | fdr_bh | 0.3688799554 |
| S3 | THb_cc | OHb_rs | spearman | two-sided | 0.6133208255 | 4.92E-05 | fdr_bh | 0.9924295604 |
| S3 | THb_cc | OHb_cf | spearman | two-sided | 0.5362101313 | 0.00056666397 | fdr_bh | 0.9573736439 |
| S3 | THb_cc | OHb_cc | spearman | two-sided | 0.8900562852 | 5.60E-13 | fdr_bh | 1 |
| S3 | THb_cc | DHb_rs | spearman | two-sided | -0.6131332083 | 4.92E-05 | fdr_bh | 0.9923909774 |
| S3 | THb_cc | DHb_cf | spearman | two-sided | -0.3609756098 | 0.02341629746 | fdr_bh | 0.6421418264 |
| S3 | THb_cc | DHb_cc | spearman | two-sided | -0.5170731707 | 0.00095192992 | fdr_bh | 0.9404305553 |
| S3 | OHb_rs | OHb_cf | spearman | two-sided | 0.6682926829 | 1.10E-05 | fdr_bh | 0.9986569814 |
| S3 | OHb_rs | OHb_cc | spearman | two-sided | 0.7857410882 | 1.37E-08 | fdr_bh | 0.9999972781 |
| S3 | OHb_rs | DHb_rs | spearman | two-sided | -0.839587242 | 1.55E-10 | fdr_bh | 0.9999999832 |
| S3 | OHb_rs | DHb_cf | spearman | two-sided | -0.6478424015 | 2.03E-05 | fdr_bh | 0.9972862017 |
| S3 | OHb_rs | DHb_cc | spearman | two-sided | -0.6628517824 | 1.14E-05 | fdr_bh | 0.9983679947 |
| S3 | OHb_cf | OHb_cc | spearman | two-sided | 0.6281425891 | 3.03E-05 | fdr_bh | 0.9950145014 |
| S3 | OHb_cf | DHb_rs | spearman | two-sided | -0.6315196998 | 2.98E-05 | fdr_bh | 0.9954880318 |
| S3 | OHb_cf | DHb_cf | spearman | two-sided | -0.5084427767 | 0.001164668965 | fdr_bh | 0.9314267042 |
| S3 | OHb_cf | DHb_cc | spearman | two-sided | -0.492120075 | 0.001677174019 | fdr_bh | 0.9119381243 |

| | | | | | | | | | | |
|---|---|---|---|---|---|---|---|---|---|---|
| S3 | OHb_cc | DHb_rs | spearman | two-sided | -0.7459662289 | 1.70E-07 | fdr_bh | 0.9999621175 | | |
| S3 | OHb_cc | DHb_cf | spearman | two-sided | -0.5968105066 | 8.24E-05 | fdr_bh | 0.9883735858 | | |
| S3 | OHb_cc | DHb_cc | spearman | two-sided | -0.6639774859 | 1.14E-05 | fdr_bh | 0.9984317428 | | |
| S3 | DHb_rs | DHb_cf | spearman | two-sided | 0.6112570356 | 5.03E-05 | fdr_bh | 0.9919963339 | | |
| S3 | DHb_rs | DHb_cc | spearman | two-sided | 0.7606003752 | 7.40E-08 | fdr_bh | 0.9999843912 | | |
| S3 | DHb_cf | DHb_cc | spearman | two-sided | 0.6300187617 | 2.98E-05 | fdr_bh | 0.9952823618 | | |
| **Figure** | **Hb** | **cond** | **test** | **tail** | **T** | **p-val** | **power** | **BF10** | | |
| S4 B | THb | cl | T-test | two-sided | -7.477467661 | 2.98E-06 | 1 | 6535.951 | | |
| S4 C | THb | cm | T-test | two-sided | -10.78919537 | 3.62E-08 | 1 | 3.57E+05 | | |
| S4 D | THb | ch | T-test | two-sided | -7.367574418 | 3.52E-06 | 1 | 5620.865 | | |
| S4 B | OHb | cl | T-test | two-sided | -6.925854779 | 7.03E-06 | 1 | 3026.086 | | |
| S4 C | OHb | cm | T-test | two-sided | -8.751777275 | 4.74E-07 | 1 | 3.43E+04 | | |
| S4 D | OHb | ch | T-test | two-sided | -6.720772115 | 9.77E-06 | 1 | 2253.917 | | |
| S4 B | DHb | cl | T-test | two-sided | 5.605146471 | 6.49E-05 | 1 | 419.525 | | |
| S4 C | DHb | cm | T-test | two-sided | 8.7177353 | 4.97E-07 | 1 | 3.29E+04 | | |
| S4 D | DHb | ch | T-test | two-sided | 7.695844802 | 2.14E-06 | 1 | 8787.149 | | |
| **Figure** | **Hb** | **Test** | **F** | **p-val** | **ng2** | **eps** | | | | |
| S5 A | THb | anova_rm | 0.142388221 | 0.836101161 | 0.002129265492 | 0.8574026346 | | | | |
| S5 A | OHb | anova_rm | 0.8575287216 | 0.4084718668 | 0.012056387 | 0.8127922358 | | | | |
| S5 A | DHb | anova_rm | 0.0492147813 | 0.9520061779 | 0.000880056189 | 0.9249617483 | | | | |
| **Figure** | **Hb** | **Test** | **F** | **p-val** | **ng2** | **eps** | | | | |
| S5 B | THb | anova_rm | 0.7901383826 | 0.4614933828 | 0.02425291708 | 0.9986208008 | | | | |
| S5 B | OHb | anova_rm | 1.674003787 | 0.2017602374 | 0.04856769248 | 0.7542993501 | | | | |
| S5 B | DHb | anova_rm | 3.968225339 | 0.02770738552 | 0.1392649898 | 0.9787464826 | | | | |
| | | **post hoc** | **A** | **B** | **T** | **Tail** | **p-val** | **corr** | **eta-square** | **BF10** |
| | | t-test | DHb_h1 | DHb_l1 | -1.730589598 | two-sided | 0.1509476579 | fdr_bh | 0.11068091 | 0.831 |
| | | t-test | DHb_h1 | DHb_l2 | -2.998867973 | two-sided | 0.0231130617 | fdr_bh | 0.16030517 | 6.467 |
| | | t-test | DHb_l1 | DHb_l2 | -0.9390763658 | two-sided | 0.3601231344 | fdr_bh | 0.02444843 | 0.35 |
| **Figure** | **Hb** | **Test** | **tail** | **T** | **p-val** | **power** | **BF10** | | | |
| S5 C | THb | T-test | two-sided | 0.3508377253 | 0.7278873157 | 0.0642876307 | 0.294 | | | |
| S5 C | OHb | T-test | two-sided | 2.001927035 | 0.05185902467 | 0.4570879881 | 1.418 | | | |
| S5 C | DHb | T-test | two-sided | -0.2732887115 | 0.7860643734 | 0.05767835824 | 0.288 | | | |
| **Figure** | **X** | **Y** | **method** | **tail** | **r** | **p-val** | **corr** | **power** | | |
| S6 | THb_ch1 | THb_cl1 | spearman | two-sided | 0.6087719298 | 0.02268653139 | fdr_bh | 0.8235262665 | | |
| S6 | THb_ch1 | THb_cl2 | spearman | two-sided | 0.1631578947 | 0.5341931162 | fdr_bh | 0.1026338979 | | |
| S6 | THb_ch1 | OHb_ch1 | spearman | two-sided | 0.7964912281 | 0.00161675269 | fdr_bh | 0.9933871752 | | |

| | | | | | | | | |
|---|---|---|---|---|---|---|---|---|
| S6 | THb_ch1 | OHb_cl1 | spearman | two-sided | 0.6561403509 | 0.0164324785 | fdr_bh | 0.8944050985 |
| S6 | THb_ch1 | OHb_cl2 | spearman | two-sided | 0.5245614035 | 0.05431644325 | fdr_bh | 0.6635326708 |
| S6 | THb_ch1 | DHb_ch1 | spearman | two-sided | -0.3210526316 | 0.2162026265 | fdr_bh | 0.2747904283 |
| S6 | THb_ch1 | DHb_cl1 | spearman | two-sided | -0.2666666667 | 0.3132791593 | fdr_bh | 0.2003626834 |
| S6 | THb_ch1 | DHb_cl2 | spearman | two-sided | -0.5157894737 | 0.05710034105 | fdr_bh | 0.6454438047 |
| S6 | THb_cl1 | THb_cl2 | spearman | two-sided | 0.5421052632 | 0.04568380188 | fdr_bh | 0.6992161172 |
| S6 | THb_cl1 | OHb_ch1 | spearman | two-sided | 0.4912280702 | 0.06923077716 | fdr_bh | 0.5943320883 |
| S6 | THb_cl1 | OHb_cl1 | spearman | two-sided | 0.6298245614 | 0.01978168859 | fdr_bh | 0.8572071211 |
| S6 | THb_cl1 | OHb_cl2 | spearman | two-sided | 0.4350877193 | 0.1186849978 | fdr_bh | 0.4786541665 |
| S6 | THb_cl1 | DHb_ch1 | spearman | two-sided | -0.4368421053 | 0.1186849978 | fdr_bh | 0.482182339 |
| S6 | THb_cl1 | DHb_cl1 | spearman | two-sided | -0.4105263158 | 0.1322708867 | fdr_bh | 0.4302182026 |
| S6 | THb_cl1 | DHb_cl2 | spearman | two-sided | -0.6754385965 | 0.0164324785 | fdr_bh | 0.9179073721 |
| S6 | THb_cl2 | OHb_ch1 | spearman | two-sided | 0.3385964912 | 0.2025171966 | fdr_bh | 0.3022070716 |
| S6 | THb_cl2 | OHb_cl1 | spearman | two-sided | 0.2 | 0.4491065601 | fdr_bh | 0.1310642523 |
| S6 | THb_cl2 | OHb_cl2 | spearman | two-sided | 0.6228070175 | 0.01978168859 | fdr_bh | 0.8463458018 |
| S6 | THb_cl2 | DHb_ch1 | spearman | two-sided | -0.2070175439 | 0.4445056859 | fdr_bh | 0.137252612 |
| S6 | THb_cl2 | DHb_cl1 | spearman | two-sided | -0.4105263158 | 0.1322708867 | fdr_bh | 0.4302182026 |
| S6 | THb_cl2 | DHb_cl2 | spearman | two-sided | -0.4 | 0.1404293427 | fdr_bh | 0.4100889069 |
| S6 | OHb_ch1 | OHb_cl1 | spearman | two-sided | 0.5964912281 | 0.02367902992 | fdr_bh | 0.8024438713 |
| S6 | OHb_ch1 | OHb_cl2 | spearman | two-sided | 0.6684210526 | 0.0164324785 | fdr_bh | 0.9097432922 |
| S6 | OHb_ch1 | DHb_ch1 | spearman | two-sided | -0.3824561404 | 0.1591267971 | fdr_bh | 0.3775155999 |
| S6 | OHb_ch1 | DHb_cl1 | spearman | two-sided | -0.0649122807 | 0.8143901624 | fdr_bh | 0.0573681965 |
| S6 | OHb_ch1 | DHb_cl2 | spearman | two-sided | -0.3701754386 | 0.1710024913 | fdr_bh | 0.355505772 |
| S6 | OHb_cl1 | OHb_cl2 | spearman | two-sided | 0.6403508772 | 0.01885484403 | fdr_bh | 0.8727727832 |
| S6 | OHb_cl1 | DHb_ch1 | spearman | two-sided | -0.6631578947 | 0.0164324785 | fdr_bh | 0.9033323178 |
| S6 | OHb_cl1 | DHb_cl1 | spearman | two-sided | -0.3333333333 | 0.2025171966 | fdr_bh | 0.2938124504 |
| S6 | OHb_cl1 | DHb_cl2 | spearman | two-sided | -0.5947368421 | 0.02367902992 | fdr_bh | 0.7993529497 |
| S6 | OHb_cl2 | DHb_ch1 | spearman | two-sided | -0.4192982456 | 0.1322708867 | fdr_bh | 0.447296266 |
| S6 | OHb_cl2 | DHb_cl1 | spearman | two-sided | -0.350877193 | 0.1949191697 | fdr_bh | 0.3223475095 |
| S6 | OHb_cl2 | DHb_cl2 | spearman | two-sided | -0.5105263158 | 0.05741120139 | fdr_bh | 0.634534799 |
| S6 | DHb_ch1 | DHb_cl1 | spearman | two-sided | 0.01052631579 | 0.9658856102 | fdr_bh | 0.0494386531 |
| S6 | DHb_ch1 | DHb_cl2 | spearman | two-sided | 0.3333333333 | 0.2025171966 | fdr_bh | 0.2938124504 |
| S6 | DHb_cl1 | DHb_cl2 | spearman | two-sided | 0.5736842105 | 0.03066774314 | fdr_bh | 0.7608717253 |
| **Figure** | **Y** | **X** | **method** | **tail** | **r** | **p-val** | **corr** | **power** |
| S7 A | AQ | THb_ch1 | spearman | two-sided | -0.2281350257 | 0.77369489 | fdr_bh | 0.1507192207 |
| S7 B | AQ | THb_cl1 | spearman | two-sided | -0.1010461073 | 0.77369489 | fdr_bh | 0.0679236948 |
| S7 A | AQ | OHb_ch1 | spearman | two-sided | -0.07291987122 | 0.77369489 | fdr_bh | 0.0588648849 |
| S7 B | AQ | OHb_cl1 | spearman | two-sided | -0.1375060429 | 0.77369489 | fdr_bh | 0.0843789224 |
| S7 A | AQ | DHb_ch1 | spearman | two-sided | 0.1958419399 | 0.77369489 | fdr_bh | 0.1226659826 |
| S7 B | AQ | DHb_cl1 | spearman | two-sided | -0.2375104377 | 0.77369489 | fdr_bh | 0.1598321171 |

| Figure | Y | X | method | tail | r | p-val | corr | power |
|---|---|---|---|---|---|---|---|---|
| S7 C | AQ | THb_ch1 | spearman | two-sided | -0.3709176048 | 0.3925638006 | fdr_bh | 0.3388451079 |
| S7 D | AQ | THb_cl1 | spearman | two-sided | -0.369869815 | 0.3925638006 | fdr_bh | 0.3370865347 |
| S7 C | AQ | OHb_ch1 | spearman | two-sided | 0.03352927501 | 0.8949238024 | fdr_bh | 0.0512206411 |
| S7 D | AQ | OHb_cl1 | spearman | two-sided | -0.2776643087 | 0.3969072558 | fdr_bh | 0.2039643689 |
| S7 C | AQ | DHb_ch1 | spearman | two-sided | 0.2179402876 | 0.4619653399 | fdr_bh | 0.1413099487 |
| S7 D | AQ | DHb_cl1 | spearman | two-sided | 0.2975723157 | 0.3969072558 | fdr_bh | 0.2289693136 |
| **Figure** | **Y** | **X** | **method** | **tail** | **r** | **p-val** | **corr** | **power** |
| S7 E | AQ | THb_ch1 | spearman | two-sided | -0.3272801904 | 0.5548063197 | fdr_bh | 0.2701568168 |
| S7 F | AQ | THb_cl1 | spearman | two-sided | -0.4561142971 | 0.3426507414 | fdr_bh | 0.49687182 |
| S7 E | AQ | OHb_ch1 | spearman | two-sided | -0.1516917708 | 0.657552151 | fdr_bh | 0.0922901976 |
| S7 F | AQ | OHb_cl1 | spearman | two-sided | -0.112210351 | 0.657552151 | fdr_bh | 0.0723840379 |
| S7 E | AQ | DHb_ch1 | spearman | two-sided | 0.1184442594 | 0.657552151 | fdr_bh | 0.0750940848 |
| S7 F | AQ | DHb_cl1 | spearman | two-sided | 0.1911731906 | 0.657552151 | fdr_bh | 0.1190276925 |
